# Regulation of fatty acid delivery to metastases by tumor endothelium

**DOI:** 10.1101/2024.04.02.587724

**Authors:** Deanna N. Edwards, Shan Wang, Wenqiang Song, Laura C. Kim, Verra M. Ngwa, Yoonha Hwang, Kevin C. Ess, Mark R. Boothby, Jin Chen

**Author notes:** Correspondence should be addressed to: Jin Chen, M.D., Ph.D., Professor of Medicine and Cell and Developmental Biology, Vanderbilt University Medical Center, T-3207D Medical Center North, 1161 21^st^ Avenue South, Nashville, TN 37232.

## Abstract

Tumor metastasis, the main cause of death in cancer patients, requires outgrowth of tumor cells after their dissemination and residence in microscopic niches. Nutrient sufficiency is a determinant of such outgrowth^1^. Fatty acids (FA) can be metabolized by cancer cells for their energetic and anabolic needs but impair the cytotoxicity of T cells in the tumor microenvironment (TME)^2, 3^, thereby supporting metastatic progression. However, despite the important role of FA in metastatic outgrowth, the regulation of intratumoral FA is poorly understood. In this report, we show that tumor endothelium actively promotes tumor growth and restricts anti-tumor cytolysis by transferring FA into developing metastatic tumors. This process uses transendothelial fatty acid transport via endosome cargo trafficking in a mechanism that requires mTORC1 activity. Thus, tumor burden was significantly reduced upon endothelial-specific targeted deletion of Raptor, a unique component of the mTORC1 complex (Rptor^ECKO^). In vivo trafficking of a fluorescent palmitic acid analog to tumor cells and T cells was reduced in Rptor^ECKO^ lung metastatic tumors, which correlated with improved markers of T cell cytotoxicity. Combination of anti-PD1 with RAD001/everolimus, at a low dose that selectively inhibits mTORC1 in endothelial cells^4^, impaired FA uptake in T cells and reduced metastatic disease, corresponding to improved anti-tumor immunity. These findings describe a novel mechanism of transendothelial fatty acid transfer into the TME during metastatic outgrowth and highlight a target for future development of therapeutic strategies.

## Background

Proliferative outgrowth of metastatic tumor cells requires sufficient nutrient availability to support energetics and biomass production^1^. Although the exact metabolic requirements for metastasis have not been fully defined, recent evidence suggests that fatty acid uptake is necessary for metastatic progression^5–7^. Compared to normal lung or primary mammary tumors, the interstitial fluid of lung metastatic tumors has drastically elevated levels of the long-chain fatty acids (LCFA) such as palmitate, serving to support the high metabolic demands of disseminated tumor cells^6^. Other studies have indicated that fatty acid enriched environments suppress cytotoxic T lymphocyte activity^2, 3^, capable of further supporting metastatic outgrowth. However, there is no indication how fatty acid delivery is regulated in the early lung metastatic niche.

The vascular system is well recognized as a critical distributor of oxygen and nutrients. Endothelial cells lining vessel walls have been described to deliver free fatty acids from the blood to surrounding tissues via a transendothelial delivery mechanism^8, 9^. The process is dependent on vascular endothelial growth factor B (VEGF-B) signaling through VEGF receptor 1 (VEGFR1) and most prominently occurs in adipose tissue, heart, and skeletal muscle^8, 9^. Like these highly metabolic tissues, tumors are excessive producers of VEGF-B^10–12^, suggesting that tumor vasculature may support delivery of fatty acids to tumor cells. In contrast to VEGF-A, VEGF-B does not appear to impact angiogenesis^13–15^, but a recent study demonstrated that it supports spontaneous metastatic colonization in the lung^12^. Due to the highly vascularized nature of the lung, transendothelial delivery of fatty acids may be particularly important to support the elevated metabolic demands of tumor cells during metastatic outgrowth, but these impacts have not been studied.

The serine/threonine kinase mTOR is a common signaling node downstream of vascular growth factor signaling^16, 17^. The mTOR kinase forms two functionally distinct complexes with shared (e.g., mTOR and mLST8) and unique components (e.g., Raptor (regulatory-associated protein of mammalian target of rapamycin) in mTORC1 and Rictor (rapamycin independent companion of target of rapamycin) in mTORC2). mTORC1 integrates environmental cues with regulation of important cellular processes, including metabolism, proliferation, and survival^16, 17^. TSC2, a component of the TSC complex, through its GTPase activating domain inactivates Rheb to the inhibit mTORC1 activity^18, 19^. While highly studied in tumor cells, the role of mTORC1 in endothelial cells is lesser known but was recently demonstrated to improve vessel integrity within solid tumors^4^. However, its role in outgrowth in the early metastatic lung niche is unknown.

In this report, we demonstrate that endothelial cell-specific deletion of Raptor (Rptor^ECKO^) reduces lung metastatic outgrowth. Loss of Raptor in endothelial cells reduces intracellular long-chain fatty acid (LCFA) content and transendothelial delivery of the fluorescent palmitate BODIPY analog to metastatic tumor cells and T lymphocytes. In contrast, loss of endothelial TSC2 (Tsc2^ECKO^) increased metastatic outgrowth and transendothelial delivery of LCFA. We demonstrate that Raptor-deficient endothelial cells exhibit deficiencies in endosome trafficking, having reduced expression of calysintenin-1 (*Clstn1*) involved in anterograde endosome transport that correlates with a mTORC1 activity gene signature in human tumor-associated endothelial cells. Indeed, knockdown of *Clstn1* reduced transendothelial transport of BODIPY in control but not Rptor-deficient endothelial cells. Reduced delivery of LCFA content in Rptor^ECKO^ metastatic tumors improved anti-tumor immune responses of T cells, suggesting that selective targeting of endothelial mTORC1 may improve the tumor microenvironment during metastatic outgrowth. Using a low-dose of the mTORC1 inhibitor everolimus (RAD001) in combination with an anti-PD1 antibody reduced BODIPY uptake in metastatic tumor cells and T cells, improving T cell responses. Taken together, our findings suggest that Raptor/mTORC1 supports transendothelial delivery of long-chain fatty acids to support metastatic outgrowth.

## Results

### Endothelial mTORC1 promotes metastatic outgrowth in the lung

To evaluate how endothelial mTORC1 impacts metastatic outgrowth in the lung, we used an inducible genetically engineered mouse model to selectively delete Raptor in endothelial cells^4^. Mice carrying floxed *Rptor* alleles (Rptor^fl/fl^, referred to as WT) were crossed with mice harboring a tamoxifen-inducible Cre recombinase (Cre^ER^) under the control of the vascular endothelial cadherin (*Cdh5*/VE-Cad) promoter (referred to as Rptor^ECKO^). To model metastatic outgrowth, we intravenously injected murine Lewis lung carcinoma cells engineered to express GFP and luciferase (LLC-GFP-luc) (Fig. 1A). To allow even seeding in the lung niche that is not influenced by endothelial Raptor status, inoculation occurred prior to tamoxifen administration and cre induction (Extended Data Fig. 1A). Raptor loss in endothelial cells (Rptor^ECKO^) significantly reduced LLC lung tumor burden based on bioluminescence imaging, lung weights, and GFP intensity (Fig. 1B-E). Similar results were obtained following intravenous inoculation of MMTV-PyMT-GFP (Extended Data Fig. 1B-C) and E0771-GFP-luc (Extended Data Fig. 1D-F) murine breast cancer cells.

**Figure 1.**
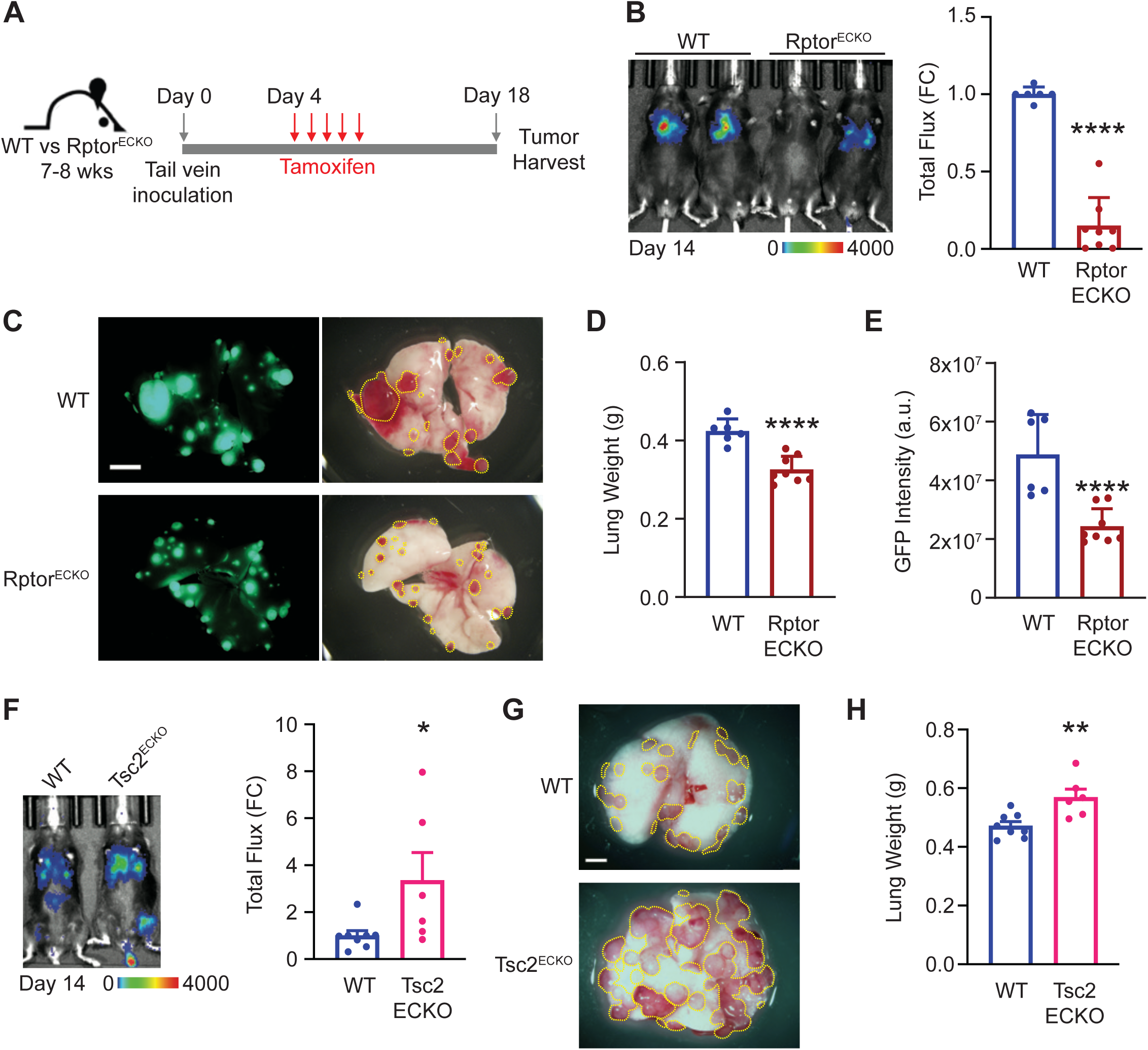
Raptor/mTORC1 loss in endothelium reduces metastatic outgrowth in the lung. **(A)** Schematic of experimental procedures of tumor cell inoculation, tamoxifen treatment, and harvest. **(B)** Representative bioluminescence image from LLC-GFP-luc tumors in WT control and Rptor^ECKO^ male mice on Day 14. Scale bar shows counts. Total radiance flux was normalized to WT controls and presented as fold change (FC). Unpaired t-test, p=1.24×10^-7^. **(C)** Representative GFP (left) and gross (right) lungs after harvest on day 18. Scale bar is 5 mm. Visible tumor area is outlined by the yellow line. **(D)** Lung weights were recorded in grams (g) at harvest and **(E)** GFP intensity was calculated as arbitrary units (a.u.). Unpaired t-test, p=1.41×10^-4^ for lung weights and p=6.69×10^-4^ for GFP intensity. **(F)** WT or Tsc2^ECKO^ female mice were inoculated with E0771-luc tumor cells as described in (A). A representative bioluminescence image is shown from Day 14. Scale bar shows counts. Total radiance flux was normalized to WT controls. Unpaired t-test, p=0.0425. **(G)** Representative lungs after harvest on Day 20 are shown. Scale bar is 5mm. Visible tumor area is outlined by the yellow line. **(H)** Lung weights were recorded at harvest. Unpaired t-test, p=0.00617. *p<0.05, **p<0.01, ***p<0.005, ****p<0.001.

To rigorously examine the role of endothelial mTORC1 on metastatic outgrowth, we also generated mice lacking endothelial TSC2 (Tsc2^ECKO^) (Extended Data Fig. 1G), a key negative regulator of mTORC1, to model increased mTORC1 activity in endothelial cells. Tsc2^ECKO^ animals developed greater tumor burden compared to controls (Fig. 1F-H), consistent with a role of endothelial mTORC1 in promoting metastatic outgrowth in the lung. However, targeting endothelial Rictor (Rictor^ECKO^), a unique and required component of mTORC2, had no significant impact on lung tumor burden (Extended Data Fig. 1H-L), indicating the critical role of endothelial mTORC1, but not mTORC2, on metastatic outgrowth in the lung.

### Long-chain fatty acids are reduced in endothelial cells upon loss of mTORC1

Due to the critical role of mTORC1 in regulating metabolism, we evaluated how mTORC1 status impacted the metabolome of control and Raptor knockout (Rptor KO) primary murine lung microvascular endothelial cells. Lipid metabolites were the most frequently altered by *Rptor* loss, particularly saturated long-chain fatty acids (LCFA) and long-chain polyunsaturated fatty acids (PUFA) (Fig.2A-B and Extended Data Fig. 2A). Specifically, LCFA and PUFA content was collectively reduced in Rptor KO endothelial cells (Fig. 2C-E), while shorter chain LCFA and medium chain fatty acid (MCFA) content was unaffected (Extended Data Fig. 2B).

**Figure 2.**
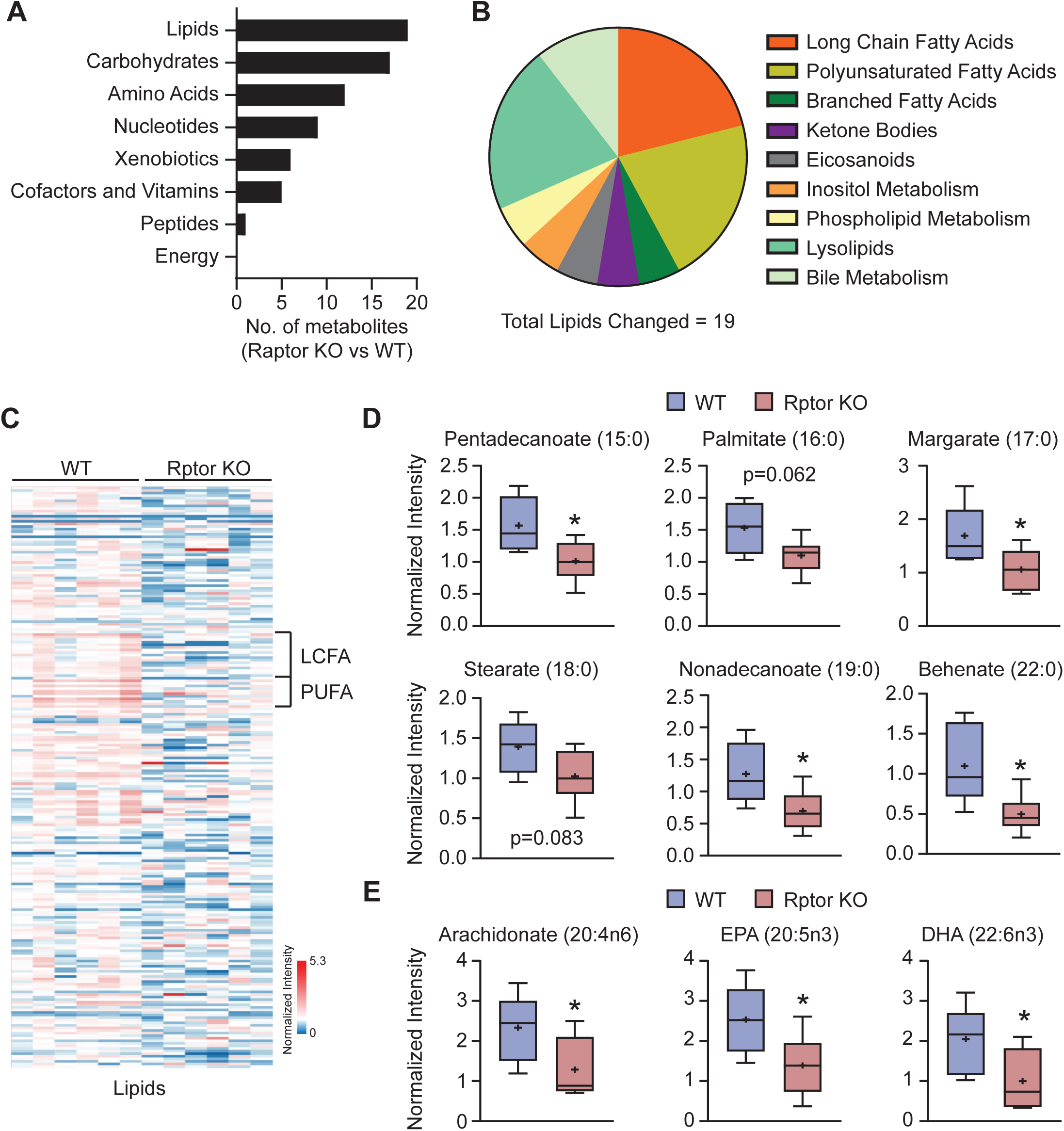
Long-chain fatty acid species are reduced upon loss of Raptor/mTORC1 in endothelial cells. Metabolomics was performed on primary microvascular endothelial cells isolated from Rptor^fl/fl^ mice transduced with control (WT) or cre-recombinase (Rptor KO) adenoviruses. **(A-B)** Summary of significantly altered metabolites by (A) major class and (B) lipid classes in Rptor KO vs WT endothelial cells. **(C)** Heat map of lipid metabolites in WT and Rptor KO endothelial cells. Columns represent individual samples and rows are metabolites. Long-chain fatty acids (LCFA) and long-chain polyunsaturated fatty acids (PUFA) are indicated. **(D-E)** Normalized intensities (log_2_+1) of representative (D) LCFA and (E) PUFA metabolites. Welch’s t-test; 15:0, p=0.0284; 16:0, p=0.0618; 17:0, p=0.0447; 18:0, p=0.0833; 19:0, p=0.0363; 22:0, p=0.0305; 20:4n6, p=0.0458; 20:5n3, p=0.0381; 22:6n3, p=0.0443. *p<0.05.

### Endothelial mTORC1 increases long-chain fatty acid content of endothelial cells downstream of VEGF-B/VEGFR1

The reduction in long-chain fatty acid content in Rptor KO endothelial cells suggests that mTORC1 regulates intracellular fatty acid levels. VEGF-B was previously reported to increase fatty acid content by endothelial cells through VEGFR1^8^, although the signaling mechanism was not defined. Therefore, we examined if mTORC1 increases fatty acid levels downstream of VEGF-B and VEGFR1 in primary lung murine microvascular endothelial cells. VEGF-B significantly increases intracellular content of BODIPY-C16, a fluorescent palmitate analog, in control but not Rptor KO endothelial cells (Extended Data Fig. 2C-E). Use of control conditions, VEGFR1 neutralizing antibody, or VEGF-A had not impact on intracellular BODIPY-C16 content, indicating that VEGF-B regulates LCFA content via mTORC1. In contrast, neither VEGF-B nor Raptor status had any impact on intracellular levels of BODIPY-C12, a medium-chain fatty acid (MCFA) laurate fluorescent analog (Extended Data Fig. 2F-H), indicating that VEGF-B/VEGFR1 and mTORC1 specifically promote LCFA content in endothelial cells.

Given that VEGF-B promotes LCFA content in a Raptor/mTORC1-dependent manner, we evaluated whether VEGF-B promotes mTORC1 activity in endothelial cells. VEGF-B increased phosphorylation of downstream mTORC1 targets, including S6K1, S6RP, and 4EBP1, but had no significant impact on phosphorylation of the mTORC2 target AKT (Extended Data Fig. 3A-B). Not surprisingly, Rptor KO endothelial cells exhibited reduced mTORC1 activity which was not impacted by VEGF-B (Extended Data Fig. 3A-B). Likewise, use of a VEGFR1 neutralizing antibody significantly reduced pS6 levels in tumor-associated endothelial cells in vivo (Extended Data Fig. 3C). Overall, these findings indicate that VEGF-B and mTORC1 augments LCFA content in endothelial cells via the same pathway.

### Endothelial mTORC1 promotes transendothelial delivery of long-chain fatty acids

In addition to its role in fatty acid levels, VEGF-B has also been demonstrated to promote transendothelial delivery of fatty acids to tissues. To examine whether Raptor/mTORC1 promotes transendothelial LCFA delivery in response to VEGF-B, we utilized a transwell assay to measure BODIPY-C16 movement across a confluent endothelial cell layer toward basolateral-located tumor cells (Fig. 3A). VEGFR1-Fc was added to remove any trace of VEGFs in the serum. Endothelial cells were carefully removed prior to imaging (Fig. 3B). With control endothelial cells, VEGF-B increases BODIPY-C16 uptake by tumor cells compared to the VEGFR1-Fc control (Fig. 3B-C). In contrast, tumor cell BODIPY-C16 uptake was unaffected by VEGF-B with Rptor KO endothelial cells (Fig. 3B-C), indicating the requirement of mTORC1 activity to support transendothelial fatty acid delivery. Endothelial permeability was not significantly impacted by VEGF-B or *Rptor* loss (Extended Data Fig. 3D-E), suggesting that LCFA are not delivered via diffusion. In agreement with a transcytosis mechanism, confocal microscopy revealed less BODIPY-C16 signal at the basolateral region of Rptor KO endothelial cells (Fig. 3D). Upon hyperactivation of mTORC1 via *Tsc2* deletion, increased transendothelial delivery of BODIPY-C16 to tumor cells was observed even in the presence of VEGFR1-Fc, while VEGF-B had no further impact (Extended Data Fig. 3F), together indicating that mTORC1 enhances transendothelial LCFA delivery through a VEGF-B dependent transcytosis mechanism.

**Figure 3.**
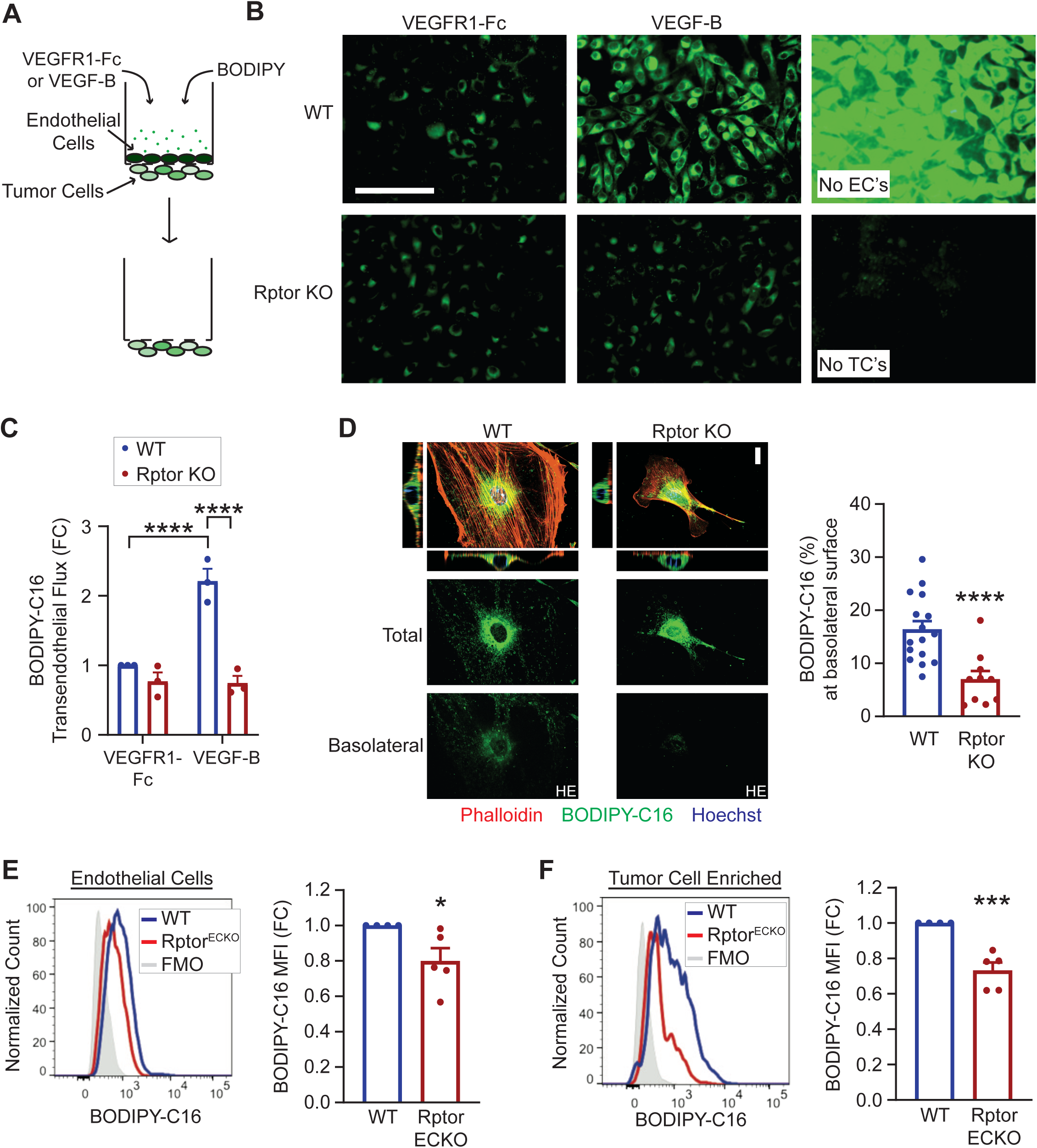
Endothelial Raptor/mTORC1 supports transendothelial delivery of FA. **(A)** Schematic of transendothelial transport assay, where BODIPY FL C16 is added to a transwell coated with a confluent top layer of WT or Rptor KO primary microvascular endothelial cells and LLC tumor cells on the bottom side. **(B)** Representative images of BODIPY FL C16 (green) in LLC tumor cells after 1 hour. Controls without endothelial cells (ECs) or tumor cells (TCs) are shown. Scale bar is 100 μm. **(C)** BODIPY-C16 fluorescence in LLC tumor cells, normalized to WT + VEGFR1-Fc control. Two-way ANOVA (p=9.73×10^-4^) with Tukey’s post hoc. **(D)** Representative spinning-disk confocal microscopy images of WT or Rptor KO primary microvascular endothelial cells treated with BODIPY FL C16 for 5 min. Nuclear (Hoechst, blue) and actin (Phalloidin, red) staining was used to detect perinuclear and cell boundaries, respectively. BODIPY-C16 (green) staining from total (all z-planes) and basolateral surface (bottom 10% of z-planes) are also shown, with the basolateral BODIPY presented at higher exposure (HE). Scale bar is 20 μm. BODIPY-C16 intensities at the basolateral surface were normalized to total signal in each cell. Unpaired t-test, p=5.38×10^-4^. **(E-F)** WT or Rptor^ECKO^ mice were inoculated with LLC tumor cells as described in Figure 1. BODIPY-C16 (50 μg) was injected 1 hr prior to tumor harvest. BODIPY-C16 median fluorescence intensity (MFI) was determined by flow cytometry in (A) CD45^-^CD31^+^ endothelial cells and (B) CD45^-^EpCAM^+^FSC^hi^ tumor-cell enriched populations and normalized to WT controls. Representative histograms are shown. Unpaired t-test; CD31^+^, p=0.0475; tumor-cell enriched, p=1.453×10^-3^. *p<0.05, **p<0.01, ***p<0.005, ****p<0.001.

To evaluate whether endothelial mTORC1 promotes LCFA delivery to tumors in vivo, we examined BODIPY-C16 uptake in lung metastatic tumors derived from WT or Rptor^ECKO^ mice. BODIPY-C16 fluorescence intensity was significantly reduced in endothelial cells from Rptor^ECKO^ mice, while Tsc2^ECKO^ tumor endothelial cells exhibited elevated uptake (Fig. 3E and Extended Data Fig. 3G). Likewise, BODIPY-C16 uptake in the tumor cell-enriched population was lower in Rptor^ECKO^ tumors but higher in Tsc2^ECKO^ tumors (Fig. 3F. and Extended Data Fig. 3H). In contrast, BODIPY-C16 uptake by tumor or endothelial cells was unaffected by endothelial-specific Rictor loss (Extended Data Fig. 3I-J), suggesting that endothelial mTORC1 regulates LCFA delivery into lung metastatic tumors.

### RAB- and CLSTN1-dependent endosome trafficking of long-chain fatty acids is defective in endothelial cells upon loss of mTORC1

RNA-seq analysis of Rptor KO lung microvascular endothelial cells revealed a downregulation of genes associated with Rab endosome trafficking (Fig. 4A-B). Immunofluorescence imaging demonstrated a significant reduction of RAB5, a marker of early endosomes, and RAB7, a marker of endosome maturation, in Rptor KO endothelial cells (Fig. 4C). Alternatively, RAB5 and RAB7 were significantly increased in TSC2 KO endothelial cells (Extended Data Fig. 4A), indicating the importance of mTORC1 in endosome trafficking in endothelial cells. Interestingly, VEGF-B exacerbates the deficiencies of RAB5 and RAB7 endosomes in Rptor KO endothelial cells (Extended Data Fig. 4B). Further transcriptome analysis of Rptor KO endothelial cells revealed a downregulation of a key mediator of anterograde vesicle transport, calsyntenin-1 (*Clstn1*), while the presence of VEGF-B exacerbates the difference in CLSTN1 protein levels^20, 21^ (Fig. 4D and Extended Data Fig. 4C-D). In contrast, TSC2 KO endothelial cells exhibit elevated *Clstn1* expression (Extended Data Fig. 4E). Together, these data suggest that Rptor KO endothelial cells exhibit defective vesicular trafficking.

**Figure 4.**
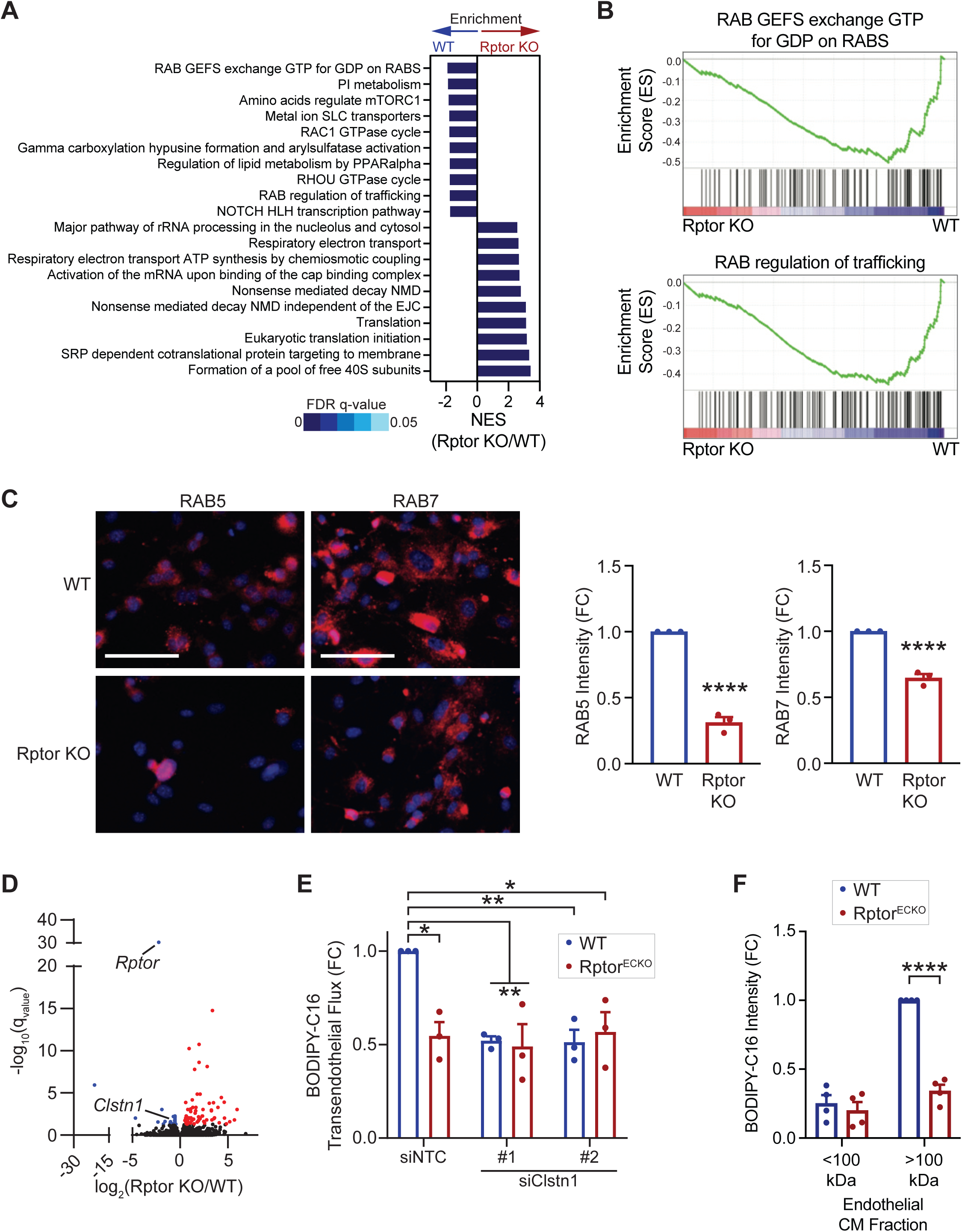
Raptor/mTORC1 loss reduces vesicle trafficking of fatty acids in endothelial cells. **(A-B)** Transcriptomic analysis of WT or Rptor KO primary microvascular endothelial cells, generated as in Figure 2. **(A)** The top gene sets enriched in WT or Rptor KO endothelial cells were determined from gene set enrichment analysis (GSEA). Normalized enrichment score (NES) and false discovery rate (FDR) q-values are indicated. **(B)** GSEA enrichment plots for RAB trafficking pathways are shown. **(C)** Immunofluorescence of RAB5 (left) or RAB7 (right) was performed on WT or Rptor KO primary microvascular endothelial cells. Representative immunofluorescence images of RAB5 and RAB 7 are shown (both red). Nuclei are stained with DAPI (blue). Scale bar is 100 μm. RAB5 or RAB7 intensities were normalized to WT controls. Unpaired t-test; RAB5, p=7.68×10^-5^; RAB7, p=3.17×10^-4^. **(D)** Volcano plot of altered gene expression in WT and Rptor KO endothelial cells from (A-B). Differentially upregulated genes in Rptor KO cells are displayed in red, while downregulated genes are in blue. **(E)** Transendothelial transport of BODIPY-C16 in WT or Rptor KO primary microvascular endothelial cells transfected with siRNA against Clstn1 was performed as described in Figure 3. BODIPY-C16 intensity in LLC tumor cells was normalized to the non-targeting control (NTC) and presented as fold change (FC). Two-way ANOVA (p=0.0137) with Tukey’s post hoc. **(F)** LLC tumor cells were incubated with the small (<100 kDa) or large (>100 kDa) fractionated conditioned media from BODIPY-C16 treated WT or Rptor KO primary microvascular endothelial cells. BODIPY intensity was normalized to the WT >100 kDa intensity. Two-way ANOVA (p=3.97×10^-5^) with Sidak’s multiple comparisons test post hoc. *p<0.05, **p<0.01, ****p<0.001.

We next examined whether changes in endosome trafficking is responsible for the defects in transendothelial LCFA delivery in Rptor KO endothelial cells. Knockdown of *Clstn1* in WT endothelial cells significantly reduced transendothelial delivery of BODIPY-C16 (Fig. 4E and Extended Data Fig. 4F-G). While BODIPY-C16 transport was significantly reduced in Rptor KO endothelial cells, *Clstn1* knockdown did not further alter BODIPY-C16 intensity in basolateral tumor cells (Fig. 4E and Extended Data Fig. 4G), suggesting that CLSTN1 functions downstream of mTORC1 to support transendothelial LCFA transport. To evaluate whether LCFAs are exported as vesicle cargo, we collected conditioned media from WT or Rptor KO endothelial cells treated with BODIPY-C16 and further fractionated it based on size. Most soluble proteins and free fatty acids are present in the <100K fraction, while larger proteins and vesicles are sequestered to the >100K fraction^22^. LLC tumor cells were then incubated with fractionated conditioned endothelial cell media and BODIPY-C16 fluorescence was determined. The majority of BODIPY-C16 was located within the >100K fraction from WT endothelial cells, but the intensity was significantly reduced in tumor cells cultured with the >100K fraction from Rptor KO endothelial cells (Fig. 4F and Extended Data Fig. 4H). Very little fluorescence was observed in the <100K fraction, suggesting that most BODIPY-C16 was contained within vesicles. Together, these data support a role for mTORC1 in driving transendothelial delivery of LCFA via a vesicle-mediated mechanism.

To understand whether mTORC1 may regulate RAB/CLSNT1-dependent endothelial trafficking in human tumors, we examined the transcriptome profiles of human tumor-associated endothelial cells in breast cancer, lung cancer, and melanoma (Extended Data Fig. 5). Expression of genes associated with mTORC1 signaling positively correlated with those involved in RAB regulation of trafficking (Extended Data Fig. 5A-C). Although CLSTN1 is most closely associated with vesicle trafficking in neurons^20, 21^, we observe expression in the endothelial cells within tumors, as well as tumor cells and fibroblasts (Extended Data Fig. 5D-G). Expression of *CLSTN1* in endothelial cells was positively associated with a higher mTORC1 expressional profile, which was stronger in tumor-associated endothelial cells (Extended Data Fig. 5H-J).

### Loss of endothelial mTORC1 reduces fatty acid uptake and improves cytotoxic activation of T lymphocytes in metastatic tumors

Accumulation of fatty acids have been associated with loss of anti-tumor functionality in CD8+ T lymphocytes^2, 3^. Therefore, we next examined whether targeting endothelial mTORC1 may reduce fatty acid uptake by T cells in metastatic tumors. WT or Rptor^ECKO^ mice bearing lung metastatic tumors were injected with BODIPY-C16 prior to harvest. Flow cytometric analysis of harvested tumors revealed a significant reduction in BODIPY-C16 intensity within CD8+ tumor infiltrating lymphocytes (TILs), but no impact on the ratio of positive cells (Fig. 5A and Extended Data Fig. 6A). Similarly, CD4+ T cells exhibited a significant decrease in BODIPY-C16 intensity, but also fewer positive cells (Extended Data Fig. 6B-C). In contrast, both CD8+ and CD4+ T cells from Tsc2^ECKO^ mice contained greater BODIPY-C16 fluorescence than controls (Extended Data Fig. 6D-E), demonstrating that endothelial mTORC1 can regulate T cell LCFA uptake.

**Figure 5.**
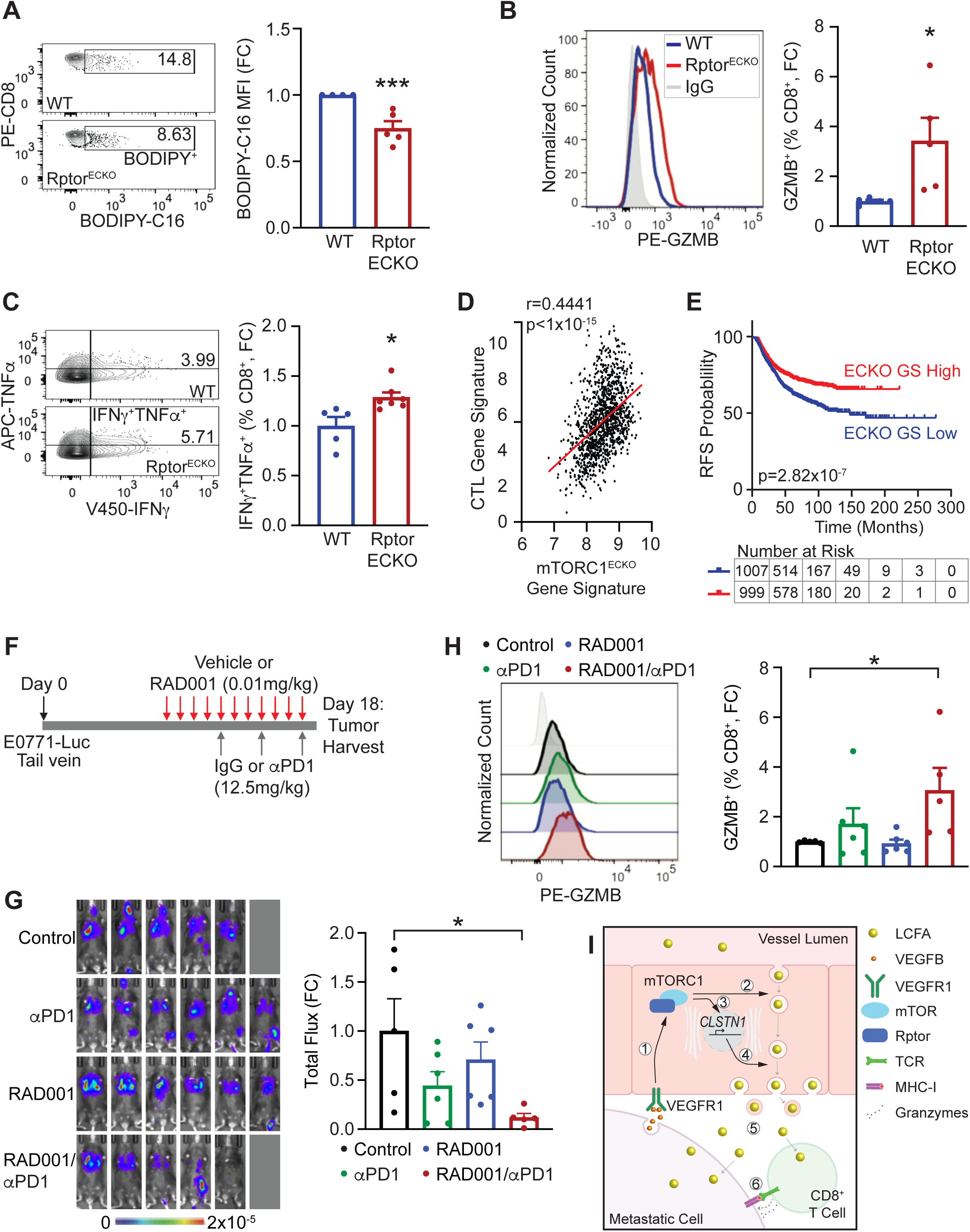
Genetic and pharmacological inhibition of endothelial mTORC1 reduces fatty acid uptake and improves anti-tumor immunity to reduce lung metastatic outgrowth. **(A)** WT or Rptor^ECKO^ male mice were inoculated with LLC tumors cells and injected with BODIPY-C16, as described in Figure 3. Representative plots of BODIPY-C16 in CD8^+^ T cells are shown. BODIPY-C16 median fluorescence intensity (MFI) was calculated and normalized to WT controls, presented as fold change (FC). Unpaired t-test, p=0.00454. **(B-C)** WT or Rptor^ECKO^ female mice were inoculated with E0771-luc cells, as described in Figure 1. Flow cytometric analysis of (B) GZMB^+^ and (C) TNFα^+^IFNγ^+^ CD8^+^ T cells was performed, and representative plots are shown. Percentages (%) of positive CD8^+^ T cells were normalized to WT controls. Unpaired t-test; GZMB^+^, p=0.0311; TNFα^+^IFNγ^+^, p=0.0128. **(D)** Correlation of the mTORC1^ECKO^ gene signature and cytotoxic T lymphocyte (CTL) gene signature in the TCGA-BRCA dataset. **(E)** Recurrence-free survival (RFS) in breast cancer patients, stratified by low (blue) or high (red) mTORC1^ECKO^ gene signature (ECKO GS). Hazard ratio (HR) is 0.6679. **(F-G)** Pharmacological mTORC1 inhibition combined with anti-PD1 immunotherapy. **(F)** Schematic of experimental procedures of tumor cell inoculation into wildtype C57BL/6 female mice, drug administration, and harvest. **(G)** Bioluminescence images from vehicle/IgG (control, black), vehicle/α PD1 (αPD1, green), RAD001/IgG (RAD001, blue), or RAD001/αPD1 (red) treated E0771-luc tumors on Day 17. Scale bar shows counts. Total radiance flux was normalized to controls and presented as fold change (FC). One-way ANOVA (p=0.0417) with Dunnett’s post hoc. **(H)** Flow cytometry was performed on harvested lung metastatic tumors for GZMB^+^ CD8^+^ T cells. Representative histogram is shown, and percentages (%) of positive CD8^+^ T cells were normalized to controls. One-way ANOVA (p=0.0475) with Dunnett’s post hoc. **(I)** Proposed model of long-chain fatty acids (LCFA) transport across endothelial cells into tumor tissue. (1) VEGFB from tumor cells activates mTORC1, leading to (2) enhanced early endosome trafficking of LCFAs via RABs. (3) mTORC1 activation also supports *CLSTN1* expression that (4) promotes anterograde transport of LCFA-filled cargos. (5) LCFAs released into the tumor tissue are taken up by metastatic cancer cells and CD8^+^ T cells, (6) resulting in reduced anti-tumor T cell responses. *p<0.05, ***p<0.005.

We next evaluated how endothelial mTORC1 impacts T cell numbers and function within metastatic tumors. Although the overall T cell composition of Rptor^ECKO^ or Tsc2^ECKO^ tumors was not significantly altered (Extended Data Fig. 6F-M)), we did observe changes in T cell function. More CD8+ T cells in Rptor^ECKO^ tumors expressed markers of cytolytic activity, including granzyme B, CD107a, and the cytokines TNFα and IFNγ (Fig. 5B-C and Extended Data Fig. 7A-C). Similarly, fewer CD8+ T cells exhibited markers of exhaustion were observed in Rptor^ECKO^ lung metastatic tumors (Extended Data Fig. 7D-E), consistent with greater functionality upon loss of endothelial mTORC1. No change was observed in cytolytic markers of CD8+ T cells from Tsc2^ECKO^ tumors (Extended Data Fig. 7F-G). In contrast to CD8+ TILs, elevated fatty acid uptake improves survival and function of T regulatory cells (Tregs)^23^. Indeed, activated Tregs characterized by high CD25 and low CD127 expression were significantly decreased in Rptor^ECKO^ tumors, but not Tsc2^ECKO^ tumors (Extended Data Fig. 7H-I). Together, these findings suggest that endothelial mTORC1 promotes fatty acid delivery to create an immunosuppressive microenvironment during metastatic outgrowth.

### Gene expression profiles of low endothelial mTORC1 activity correlate with improved cytotoxicity and progression-free survival in human cancer

To evaluate how endothelial-specific loss of Rptor impacts the tumor microenvironment, we performed RNA-seq on sorted GFP+ tumor cells from WT or Rptor^ECKO^ metastatic tumors (Extended Data Fig. 8A-D). Interestingly, we observed that Rptor^ECKO^ tumor cells exhibit reduced expression of genes associated with respiratory electron transport chain pathways but an enrichment of those in inflammatory signaling pathways (Extended Data Fig. 8D). These transcriptional changes were used to evaluate the relationship between endothelial mTORC1 activity and cytotoxic T lymphocyte (CTL) responses and survival from publicly-available bulk patient samples. The mTORC1^ECKO^ gene signature reflects the expression profile of tumor cells that are associated with low endothelial mTORC1 activity. Indeed, a CTL gene signature^24^ is positively correlated with the mTORC1^ECKO^ gene signature in breast cancer, lung cancer, and melanoma datasets (Fig. 5D and Extended Data Fig. 8E-F). Similar findings were observed in metastatic cancer datasets from breast cancer and melanoma (Extended Data Fig. 8G-H). Patients with tumors exhibiting transcriptional profiles similar to mTORC1^ECKO^ tumors also incur significantly reduced recurrence-free survival (RFS) probabilities and improved responses to pembrolizumab therapy (Fig. 5E and Extended Data Fig. 8I-J). These data reflect improved anti-tumor immune responses and survival in patients with low endothelial mTORC1 activity.

### Low-dose RAD001 in combination with anti-PD1 therapy reduces fatty acid delivery and metastatic tumor burden

The mTORC1 inhibitor, everolimus/RAD001, has been long considered to be immunosuppressive. However, recent studies have demonstrated that low doses can inhibit mTORC1 signaling without causing immunosuppression^4, 25, 26^. Further, we previously demonstrated that low-dose RAD001 selectively targets mTORC1 in endothelial cells^4^. Patient data also suggested that reduced endothelial mTORC1 activity may improve pembrolizumab/PD-1 inhibition (Extended Data Fig. 8J). Therefore, we evaluated whether low-dose RAD001 in combination with anti-PD-1 therapy would reduce lung metastatic outgrowth (Fig. 5F). Compared to vehicle/IgG control, lung tumor burden was significantly reduced in animals treated with the RAD001/αPD1 combination therapy (Fig. 5G and Extended Data Fig. 9A-C). RAD001/αPD1 significantly reduced BODIPY-C16 fluorescence in both endothelial and tumor-enriched cell populations, while a similar decrease was observed in endothelial cells of RAD001-treated mice (Extended Data Fig. 9D-E), consistent with less LCFA transport into metastatic tumors upon mTORC1 inhibition. Likewise, the combination of RAD001 with αPD1, but not either agent alone, significantly reduced BODIPY-C16 uptake by CD4+ and CD8+ T cells, while increasing T cell enrichment and cytolytic GZMB+ CTLs (Fig. 5H and Extended Data Fig. 9F-J). Together, these results suggest that endothelial mTORC1 can be therapeutically targeted to improve anti-tumor responses in combination with anti-PD-1therapy to treat metastatic progression (Fig. 5I).

## Discussion

The changing nutrient composition within the early metastatic microenvironment is a critical factor in metastatic progression. Recent studies have demonstrated the importance of fatty acids in metastasis, including their enrichment in the lung metastatic microenvironment^5–7^. However, how fatty acids are delivered to early metastatic tumors is poorly understood. The endothelium serves as a gatekeeper between the nutrient-rich blood and highly metabolic tissues. Here, we report that endothelial mTORC1 is a key driver of transendothelial fatty acid delivery to early metastatic tumors, leading to reduced anti-tumor activities by T cells to support metastatic outgrowth (Fig. 5I). Selective pharmacological inhibition of endothelial mTORC1 using a low dose of everolimus/RAD001 enhanced the efficacy of PD-1 inhibition that reduced metastatic outgrowth, with a concomitant decrease in long-chain fatty acid (LCFA) enrichment and elevated cytolytic capacity in T cells. These findings improve our understanding of how fatty acids composition is regulated in the early metastatic microenvironment and demonstrate that selective endothelial mTORC1 inhibition may improve immune checkpoint therapy responses in metastatic disease.

Long-chain fatty acids have been found to accumulate during tumor progression, including at the metastatic site^2, 6^. Multiple sources of fatty acids within tumors have been described, including stromal cell release and de novo synthesis within tumor cells^6, 27^. However, several groups have linked elevated dietary intake of lipids with cancer progression^6, 28, 29^, suggesting that the blood may be a significant source of free fatty acids in tumors. Endothelial cells that line blood vessels serve as gatekeepers to regulate dispersal of nutrients and other molecules into surrounding tissues via transendothelial delivery^8, 9^. Indeed, our data demonstrated that endothelial cells are responsible for delivery of long-chain fatty acids into the tumor microenvironment through an mTORC1-dependent mechanism. Using a murine model of metastatic outgrowth, loss of the mTORC1 component Rptor in endothelial cells reduced uptake of the fluorescent palmitate analog BODIPY-C16 within tumors that was associated with a decrease in metastatic tumor burden. These findings support fatty acid delivery from the blood as a significant source during metastatic progression within the lung. This mechanism may be a substantial source of fatty acids for disseminated tumor cells within the lung, which is a highly vascular. While a similar mechanism may also contribute to other metastatic sites, such as bone and brain, it is unclear whether specialized fenestrated sinusoidal endothelial cells within the liver, which serve to readily diffuse substrates from the blood, would undergo vesicular transport of fatty acids.

While tumor angiogenesis through VEGF-A and VEGFR2 has been extensively studied, the role of VEGF-B and its cognate receptor VEGFR1 is less understood. In normal but highly metabolic tissues, VEGF-B has been described as a driver of transendothelial fatty acid delivery^8, 9^, although the signaling mechanism has not been defined. Our findings describe a critical role of mTORC1 in VEGF-B-associated transport of long-chain fatty acids in endothelial cells. While VEGFR1 is necessary for VEGF-B activation of mTORC1, neuropilin-1 (NRP1) is also required for fatty acid transport across endothelial cells but it is unclear how this co-receptor may be involved in the signaling cascade^8, 30–33^. Like skeletal muscle, heart and adipose tissues, VEGF-B expression is elevated in human disease, including increased expression within tumors and cell lines^10–12, 34^. Still, its role in cancer has been largely unknown, which may be somewhat attributed to both pro- and anti-angiogenic properties described for VEGF-B^13–15^. It remains possible the VEGF-B isoforms, including the soluble VEGF-B_186_ and membrane-bound VEGF-B_167_, may have different properties. Most studies have focused on the shorter isoform, which is more broadly expressed^10^. However, we evaluated VEGF-B_186_, which is more highly expressed in tumors and cancer cell lines and was found to have a more profound impact on fatty acid content in endothelial cells^8^, supporting the notion of a differential isoform effect. Although VEGF-B does not appear to significantly impact primary tumor growth, a study that evaluated the full-length gene found that VEGF-B does support the development of metastasis in the lungs^12^. Our data is consistent with a role of VEGF-B in promoting early metastatic outgrowth by creating a fatty acid-rich environment within the early lung metastatic. Interestingly, Yang and colleagues attributed the VEGF-B-associated metastatic spread to a decline in vessel integrity^12^. We previously demonstrated that Raptor loss in endothelial cells improves tumor vessel structure and reduces metastasis^4^. Although additional studies will be necessary to elucidate the mechanism of mTORC1 in vessel integrity, it is intriguing to speculate that changes in VEGF-B signaling or fatty acid content may contribute to mTORC1’s role in vessel function.

Endocytic trafficking across the endothelial cell lining supports delivery of nutrients to surrounding tissues, although the mechanisms regulating this process have not been thoroughly explored. Most studies have investigated those impacting the blood-brain barrier, where PTEN, a negative regulator of mTORC1 signaling, was identified as a suppressor of endothelial transcytosis^35^. Our data demonstrates that mTORC1 promotes endosomal transcytosis in microvascular lung endothelial cells to mediate transendothelial fatty acid delivery during metastatic outgrowth. Indeed, endothelial transcytosis involves RAB7 late endosomes, feeding into microvesicular bodies that release small extracellular vesicles, both of which contain palmitate as cargo^36–38^. We also demonstrated that mTORC1 regulated expression of CLSTN1, a protein implicated in endosomal trafficking and exocytosis^20, 21^. Early studies attributed the mechanism to increased fatty acid uptake, given that VEGF-B promoted expression of the fatty acid transporters FATP3 and FATP4^8^. Recent studies have suggested that FATP3 and FATP4 may serve to sequester fatty acids, particularly along the mitochondria and endoplasmic reticulum, respectively, where these proteins are localized^39–42^. This localization has significant implications for endothelial metabolism, especially given the acyl-coA synthetase activity of the FATP family^42^. We observe intense perinuclear BODIPY-C16 localization in endothelial cells, although this compartmentalization does not appear to be significantly impacted by mTORC1 status. It remains possible that autophagy activation in the absence of mTORC1 may maintain fatty acid uptake^43^, but the overall reduction in long-chain fatty acids in Raptor-deficient endothelial cells may have secondary impacts on endothelial cell fatty acid metabolism. In addition to its role in lipid transport, VEGF-B and mTORC1 has been shown to have secondary impacts on general endothelial sugar and amino acid metabolism^8, 44, 45^ that may have significant impacts on vessel structure and function in tumors^46–48^. Additional studies are necessary to more completely understand whether endothelial metabolism may contribute to endothelial function in response to targeting mTORC1.

In addition to providing fatty acids to serve as fuel for proliferating tumor cells, a growing body of evidence suggests that accumulating fatty acids within tumors has significant negative impacts on anti-tumor immunity. Although CD8+ TILs do utilize LCFAs to support effector function in some environments, those highly enriched with fatty acids lead to dysfunction^2^. Within environments enriched for LCFAs, Manzo and colleagues demonstrated that CD8+ T cells underwent transcriptional alterations that impairs their ability to utilize and safely store fatty acids^2^. The presence of palmitate reduced the ability of CD8+ T cells to produce effector cytokines and cytolytic enzymes, including IFNγ, TNFα, and Granzyme B^2^. Excessive LCFA uptake also promotes lipid peroxidation and ferroptosis in CD8+ T cells, further limiting their anti-tumor responsiveness^49, 50^. Further potentiating an immunosuppressive microenvironment, high fatty acid levels are supportive of T regulatory (Treg) cell survival^51^. In agreement, our findings demonstrated that loss of Raptor/mTORC1 in endothelial cells reduced fatty acid uptake by TILs, improved anti-tumor CD8+ T cell effector function, and reduced immunosuppressive Tregs during metastatic outgrowth. Interestingly, these improvements in anti-tumor lymphocyte responses may further enhance tumor cell death mediated by ferroptosis^52^. Additional immunosuppressive influences of accumulating fatty acids within the diverse tumor microenvironment have been described in myeloid populations that may have further implications for T cell responses. Within palmitate-enriched conditions, dendritic cells exhibit impaired antigen presentations, while tumor-associated macrophages undergo a metabolic switch toward fatty acid oxidation to support M2 polarization^53–57^. Thus, reducing fatty acid accumulation within the tumor microenvironment may help support improve anti-tumor immune responses.

Metastatic cancer remains uncurable, but recent advances in immunotherapy have offered improved outcomes in some patients. Unfortunately, the overall response rate remains low, particularly those with tumors considered immunologically cold^58^. In recent years, vascular-targeting strategies, such as anti-angiogenics, have been shown to improve immunotherapy responses in both pre-clinical models and advanced cancer patients^63–68^. As a major contributor to immune suppression, the metabolic tumor microenvironment has also been an area of intense research to identify new targets to improve immunotherapy responses^59–62^. Our findings suggest that fatty acid accumulation within metastatic tumor lesions can be reversed by targeting endothelial mTORC1 to improve immunotherapy efficacy. ^59–62^ We demonstrate that selective pharmacological inhibition of endothelial mTORC1 using low doses of everolimus/RAD001 improved anti-PD-1 responses that correlated with reduced LCFA uptake by T lymphocytes. Compared to either drug alone, the combination of low-dose RAD001 with anti-PD-1 showed the strongest impacts on T cells, reducing fatty acid uptake and improving cytolytic capacity. PD-1 signaling is also associated with a metabolic switch in CD8+ T cells toward fatty acid uptake and metabolism, leading to mitochondrial damage and ferroptosis in lipid-enriched environments^69, 70^. While a reduction in T cell BODIPY uptake appears to be partially attributed to pharmacological targeting of endothelial mTORC1, our data suggest that anti-PD1 therapy further improves the metabolic fitness of TILs that supports further improvements of anti-tumor immunity in lung metastatic tumors. Therefore, our findings suggest that vascular-targeting strategies to reduce environmental fatty acid accumulation may represent a new approach to enhancing immunotherapy responses in metastatic cancer.

## Methods

### Cell Culture

LLC, E0771, and MMTV-PyMT parental cells were maintained as previously described^4, 24, 46^. GFP and luciferase were cloned into the pCDH-puro plasmid. The lentiviral vector pLX311-luciferase was a gift from William Hahn (Addgene plasmid #117735). Parental cells were transduced with pCDH-GFP-luc or pLX311-luc. GFP^+^ cells were enriched using cell sorting. Murine pulmonary microvascular endothelial cells (MPMECs) were isolated from 8 to 16-week-old *Rptor*^fl/fl^ or *Tsc2*^fl/fl^ mice and maintained in EGM-2 medium (Lonza), as previously described^71–75^. For adenoviral Cre expression, MPMECs were seeded onto tissue culture plates coated with 0.1% gelatin at 70-80% confluency. Cells were infected with 10^7^ PFU ml^-1^ of Ad-CMV-iCre (Vector Biolabs, #1045) for 16-48 hours, as indicated. Ad-CMV-b-Gal (Vector Biolabs #1080) or Ad-CMV-Null (Vector Biolabs #1300) were used as control vectors as indicated. Cells were harvested 48-72 hours after transduction.

### Animals

All mice used in this study were maintained on a C57BL/6 background and housed in a non-barrier facility. Wildtype C57BL/6 female mice were purchased from Jackson Laboratories. CDH5-CreER^T2^ mice and animals with *Rptor* or *Rictor* floxed alleles were sourced and maintained as described previously^4, 71^. *Tsc2* floxed mice were generated and provided by Kevin Ess (Vanderbilt University Medical Center^76^. Animals were genotyped for Cre or floxed alleles of *Rptor*, *Rictor*, or *Tsc2* alleles using the primers listed in Extended Data Table 1. To induce endothelial-specific deletion of *Rptor*, *Rictor*, or *Tsc2*, tamoxifen (T5648, Sigma) was reconstituted in sunflower seed oil (S5007, Sigma) and administered intraperitoneally (i.p.) for five consecutive days in 7 to 12-week-old mice at a final dose of 2 mg/mouse, as previously described^4^. Tamoxifen was administered beginning four days after intravenous tumor cell inoculation.

### Tumor Models

For modeling metastatic outgrowth, 1×10^6^ cancer cells in 200 μL PBS were intravenously injected into the tail vein. For VEGFR1 neutralization in vivo, animals were treated with 2.5 mg/kg of normal rabbit IgG (R&D Systems #AB-108-C) or anti-VEGFR1 (R&D Systems #AF471) every 3-4 days by i.p. injection. Lung metastatic tumors were tracked weekly on live animals with bioluminescence imaging (IVIS Spectrum, PerkinElmer, Shelton, CT). Animals were sacrificed at day 14-18, and lungs were perfused with PBS through the cardiac left ventricle. Lung weights were recorded, and gross lungs imaged using an Olympus stereo microscope. GFP intensity was measured using ImageJ software as arbitrary units (a.u.), using the mean intensity from front and back side of each lung. Total radiance flux (p/sec/cm^2^/st; photons per second within a cm^2^ tissue area per steradian) was calculated in the thorax and normalized to WT littermate controls using Living Image v.4.8.2 (Perkin Elmer, Shelton, CT) or Aura v4.0.8 (Spectral Images, Tuscon, AZ) software. To assess fatty acid uptake of tumor cell populations, animals were inoculated and induced with parental or luciferase-expressing tumor cells as described above. Beginning 1 hour prior to sacrifice and harvest, animals were injected with 50 μg of BODIPY FL C16 (BODIPY-C16, Thermo #D3821) in 50 μL DMSO by i.p. injection.

For drug studies with RAD001/everolimus and anti-PD1, E0771-luc cells were injected into 8-week-old wildtype C57BL/6 female mice, as described above. Seven days after tumor inoculation, mice were treated with vehicle+IgG, vehicle+αPD1, RAD001+IgG, or RAD001+αPD1. Vehicle (20% DMSO in PBS) or 0.01 mg/kg RAD001 (Selleck Chemical, #S1120) was administered daily in 100 μL. 250 μg of IgG (BioXCell #BE0089) or anti-PD1 (clone RMP1-14, BioXCell #BE0146) was administered by i.p. in 100 μL PBS every three days beginning on day 11. Bioluminescent imaging and BODIPY FL C16 treatment was performed as above. Sections from formalin-fixed, paraffin-embedded lungs were stained with hematoxylin and eosin, as previously described^24^. Images were obtained using an Olympus upright compound microscope.

### Metabolite Profiling

MPMECs isolated from *Rptor*^fl/fl^ mice were transduced with LacZ or Cre-expressing adenovirus as described above. Cell pellets were profiled by Metabolon (Durham, NC) using ultrahigh performance LC-MS/MS to detect 541 metabolites. Compounds were identified against a library of standards, and abundance was normalized to protein content. Scaled results were grouped in major classes and subclasses. Six samples from independent MPMEC isolations were analyzed in each group.

### Flow cytometry

Lung metastatic tumors were dissociated in RPMI-1640, 5% FBS, collagenase IA (1 mg/mL, Sigma #C9891), and DNaseI (0.25 mg/mL, Sigma #DN25), filtered through a 70 μm cell strainer, and red blood cells were lysed, as previously described^24^. For detection of IFNγ and TNFα, 2×10^6^ live cells were stimulated in RPMI-1640 supplemented with 5% FBS, phorbol 12-myristate 13-acetate (PMA) (50 ng/mL, Sigma #P8139), ionomycin (1 μg/mL, Sigma #I0634), and GolgiStop protein transport inhibitor (1:1500, BD Biosciences #554724) at 37°C for 4 hrs. To exclude dead cells from analysis, Ghost Dye Violet 510 (Tonbo Biosciences #13-0870). Blocking with anti-CD16/32 (Tonbo #70-0161) was followed by staining of the following extracellular proteins with antibodies shown in Extended Data Table 2. Cytofix/Cytoperm solution kit (BD Biosciences, #554714) was used to the detect intracellular cytokines GZMB, IFNγ, or TNFα, per manufacturer’s instructions. The FoxP3/Transcription Factors Staining Kit (Tonbo #TNB-0607) was used to detect FoxP3, as directed. BODIPY-C16 was detected using the Alexa Fluor 488 channel. To detect pS6, 4×10^6^ cells were fixed in methanol and incubated with anti-pS6 (Ser235/236) rabbit antibody (1:100, Cell Signaling Technology #2211), followed by incubation with cell surface markers and secondary Alexa Fluor 647 goat anti-rabbit (1:200, Invitrogen #A21244) as described previously ^4^. Tumor cell suspensions or isolated splenocytes were used for compensation, unstained, fluorescence minus one (FMO), and isotype controls, where appropriate. Using BD FACS Diva software, flow cytometry data was obtained on a BD Fortessa, and analysis was completed using FlowJo software (v10).

### Immunoblotting

MPMECs were isolated from *Rptor*^fl/fl^ mice and transduced with Cre adenovirus for 16-48 hrs. Ad-CMV-Null was used as a control. To assess VEGF-B activation of mTORC1, cells were serum starved in EBM-2 medium supplemented with 0.2% FBS overnight. A time course was performed on cells treated with 300 ng/mL of recombinant mouse VEGF-B_186_ (R&D Systems #767-VE-010) for 0, 5, 15, or 30 minutes prior to harvest. In a separate experiment, cells were incubated with VEGF-B_186_ (0-300 ng/mL) for 30 min before harvest. Pre-cleared lysates were electrophoresed on a SDS-polyacrylamide gel and transferred to nitrocellulose membranes, as previously described^24^. The following primary antibodies were used at 1:1000 dilution: Raptor (Cell Signaling Technology (CST) #2280), Rictor (Bethyl #A300-459A), Rictor (Millipore #05-1471), phospho-S6K1 (T389, CST #9234), S6K1 (CST #9202), phospho-S6RP (S235/236, CST #2211), S6RP (CST #2217), phospho-4EBP1 (T37/46, CST #9459), phospho-4EBP1 (T70, CST #9455), 4EBP1 (CST #9644), phospho-AKT (T308, CST #4056), phospho-AKT (S473, CST #4060), AKT (CST #2920), CLSTN1 (Abcam #ab134130), and β-tubulin (Sigma #T4026). Blots were incubated with secondary antibodies IRDye 680LT goat anti-mouse (1:20,000; LI-COR #925-68020) or IRDye 800 CW goat anti-rabbit (1:10,000; LI-COR #925-32211) and imaged using LI-COR Odyssey.

### Transendothelial Delivery Assay

LLC parental cells (5×10^4^) were initially plated onto the bottom side of a 0.4 μm pore size transwell insert. After LLC attachment, 1×10^5^ *Rptor*^fl/fl^ or Tsc2^fl/fl^ MPMECs were plated into the inner side coated with 0.1% gelatin. Cells were transduced with Ad-CMV-Null or Ad-CMV-iCre for 16 hours, followed by stimulation with VEGF-B_186_ (300 ng/mL) for 30 hr in EBM-2 supplemented with 1% BSA free of fatty acids (FFA-BSA). Control cells were incubated with soluble hVEGFR1 (VEGFR1-Fc, 1 μg/mL; R&D Systems #321-FL-050). BODIPY FL C16 (1 μM) was added to the top chamber for 1 hr, and cells were thoroughly washed 3x with 1% FFA-BSA in EBM-2. Cells were fixed with 4% paraformaldehyde (PFA) for 10 min and endothelial cells were carefully removed using a cotton tip applicator. The transwell lining with tumor cells was imaged using an Olympus compound microscope. BODIPY-C16 intensity was quantitated in ImageJ and normalized to WT+VEGFR1-Fc controls.

To evaluate transport after *Clstn1* knockdown, *Rptor*^fl/fl^ MPMECs were transduced with empty or cre-expressing adenovirus for 16 hours, followed by transfection with 40 nM of ON-TARGETplus siClsnt1 (Horizon Dharmacon #J-044659-09 and #J-044659-10) or non-targeting control (Horizon Dharmacon D001810-10-20) using Lipofectamine RNAiMAX, per manufacturer’s instructions. After 24 hours, LLC and MPMECs were plated onto transwells and assayed as described above.

### Permeability Assay

1×10^5^ *Rptor*^fl/fl^ MPMECs were plated onto the inner surface of a gelatin-coated transwell insert with a 0.4 μm pore size. Cells were transduced with Ad-CMV-Null or Ad-CMV-iCre for 16 hours, and stimulated with VEGF-B_186_ (300 ng/mL) or VEGF-A (300 ng/mL) for 30 hr in EBM-2 supplemented with 1% BSA free of fatty acids (FFA-BSA). 70 kDa Texas Red-Dextran (1 mg/mL, Thermo #D1830) was placed in the lower chamber. Ten microliter aliquots of medium were removed from the upper chamber at 0, 10, 30, and 60 min. A transwell without endothelial cells was used to determine maximum permeability intensity. A BioTek Synergy HT plate reader (595 excitation, 615 emission) was used to measure fluorescence intensity of Texas Red-Dextran that passed into the upper chamber. Permeability was calculated as a percent of the empty transwell control.

### BODIPY Uptake Assay

*Rptor*^fl/fl^ MPMECs were transduced with Ad-CMV-Null or Ad-CMV-iCre for 24 hrs. Cells were stimulated with recombinant mouse VEGF-A (300 ng/mL, R&D Systems #493-MV-005), recombinant mouse VEGF-B_186_ (300 ng/mL), mouse VEGFR1/Flt-1 antibody (1 μg/mL, R&D Systems, #AF471), or a combination of anti-VEGFR1 and VEGF-B for 30 hrs. For the combination, cells were pre-incubated with anti-VEGFR1 for 2 hr prior to addition of VEGF-B. Cells were then incubated with 10 μM of BODIPY FL C16 or BODIPY 500/510 C1,C12 (BODIPY-C12, Thermo #D3823) for 3 min. For imaging, cells were fixed in 4% PFA for 10 min and stained with DAPI (Invitrogen #R37606) as directed. Cells were imaged using an Olympus inverted microscope. For flow cytometry, cells were trypsinized and fixed in 4% PFA for 10 min. Flow cytometry was immediately performed as described above.

### BODIPY Confocal Imaging

*Rptor*^fl/fl^ MPMECs were transduced with Ad-CMV-Null or Ad-CMV-iCre for 24 hrs on MatTek dishes with a No. 1.5 coverslip coated with 0.1% gelatin. Cells were stimulated with VEGF-B (300 ng/mL) for 30 hrs in EBM-2 supplemented with 1% FFA-BSA. Prior to imaging, live cells were stained with Phalloidin-iFluor594 (Abcam #ab176757) and Hoescht 33342 (Invitrogen #R37605), according to manufacturer’s instructions. Cells were incubated with BODIPY FL C16 (20 μM) for 20 min at 4°C, then washed 3x in EBM-2 with FFA-BSA. Cells were then incubated at 37°C for 5 min and immediately imaged using a Nikon Spinning Disk Microscope. Images from three regions were collected from three independent experiments. The z-plane representing the apical (top) and basolateral (bottom) edges of the cell were determined by phalloidin staining. The basolateral surface represents the bottom 10% of z-plane images for each cell. BODIPY-C16 intensity was quantitated in ImageJ, and the intensity present at the basolateral surface was calculated as a percent of whole cell intensity.

### Conditioned Medium Assay

*Rptor*^fl/fl^ MPMECs were transduced with Ad-CMV-Null or Ad-CMV-iCre for 16 hrs, then stimulated with VEGF-B (300 ng/mL) in EBM-2 supplemented with 1% FFA-BSA for 30 hrs. Endothelial cells were incubated with 5 μM BODIPY FL C16 for 1 hr and subsequently thoroughly washed to remove free BODIPY. Cells were cultured in EBM-2 + 1% FFA-BSA for 16 hr. Conditioned medium was collected and fractionated using 100 kDa spin columns (Amicon #UFC510024), and resulting fractions were re-constituted to equivalent volumes. Complete or fractionated (<100 kDa or >100 kDa) conditioned medium was added to LLC tumor cells for 4 hr. Cells were washed thoroughly and images were captured from 3 fields of view using an Olympus microscope. BODIPY-C16 intensity was calculated using ImageJ and normalized to samples cultured in medium from >100 kDa fraction from WT endothelial cells.

### Immunofluorescence

2.5×10^4^ *Rptor*^fl/fl^ or *Tsc2*^fl/fl^ MPMECs were seeded onto a 96 well plate coated with 0.1% gelatin and transduced with Ad-CMV-Null and Ad-CMV-iCre for 48 hrs. Cells were fixed with 2% PFA and permeabilized with 1% Triton X-100. Cells were blocked with 3% BSA and probed with antibodies against TSC2 (CST #4308), RAB5 (CST #3547), or RAB7 (CST #9367) using a 1:100 dilution overnight at 4°C. Cells were then incubated with Alexa Fluor 488 (Invitrogen #A11034) or Alexa Fluor 594 (Invitrogen #A11012) anti-rabbit secondary antibodies, both 1:500 for 1 hr. Nuclei were stained with DAPI, as indicated. Three fields of view per sample were obtained using an Olympus inverted fluorescent microscope. Signal intensity was quantitated using ImageJ and normalized to the appropriate control. For VEGF-B stimulation, *Rptor*^fl/fl^ MPMECs were transduced as described above for 24 hrs. Cells were then incubated with 1 μg/mL VEGFR1-Fc or 300 ng/mL VEGF-B for 30 hrs in 1% FFA-BSA prior to immunofluorescence.

### CLSTN1 Immunohistochemistry Images

CLSTN1 immunohistochemistry (Gene #ENSG00000171603; Antibody #HPA077705) images in breast lobular carcinoma^77^ (Patient ID #2900), lung squamous cell carcinoma^78^ (Patient ID #4488), and melanoma^79^ (Patient ID #4229) were obtained from The Human Protein Atlas^80^ (www.proteinatlas.org).

### RNA Sequencing

*Rptor*^fl/fl^ MPMECs were transduced with Ad-CMV-Null or Ad-CMV-iCre for 48 hrs. RNA was extracted using Trizol (Life Technologies #15596026) and RNeasy Kit (Qiagen #74104), as directed. To isolate RNA from lung metastatic tumor cells, WT or Rptor^ECKO^ male mice were inoculated with LLC-GFP-luc cells. Dissociated tumor cells were separated based on CD45 using CD45 mouse microbeads (Miltenyi Biotec, #130-052-301), according to manufacturer’s instructions. GFP+ tumor cells were collected by flow sorting, and RNA was extracted using RNAqueous Micro Total RNA Isolation Kit (Invitrogen #AM1931), according to manufacturer’s instructions. RNAseq was performed by BGI Americas (Cambridge, MA) using the DNBSEQ platform. After sequencing, raw data was filtered to remove reads with high rates of unknown bases, low quality reads, and reads of adapter sequences. Clean reads were aligned to the reference genome (*Mus musculus*, version GCF_000001635.26_GRCm38.p6) using HISAT and aligned to reference genes using BowTie2. Differentially expressed genes (DEG) were identified using DESeq2 (q value < 0.05) using the Dr. Tom platform (BGI). Pathway enrichment analysis was performed against REACTOME gene sets using Gene Set Enrichment Analysis (GSEA) Software (v4.3.2, Broad Institute)^81, 82^.

### Quantitative Real-Time PCR

RNA was collected from *Rptor*^fl/fl^ or *Tsc2*^fl/fl^ MPMECs transduced with Ad-CMV-Null or Ad-CMV-iCre, as described above. Following the manufacturer’s directions, cDNA was generated using the iScript cDNA synthesis kit (Bio-Rad #1708891). Using the primers defined in Extended Data Table 3, PCR amplification was performed in triplicate using the StepOnePlus (Applied Biosystems) as previously described^24^. The ΔΔC_t_ quantitation method was used.

### RNA Human Dataset Analysis

RNA expression data from the TCGA Breast Adenocarcinoma (BRCA v.12-17-2021, n=1218), TCGA lung cancer (LUNG v.05-26-2021, n=1129), and TCGA Melanoma (SKCM v.11-02-2022, n=474) was acquired from the UCSC Xena platform^83^ (https://xena.ucsc.edu). RNA expression data of metastatic biopsies from The Metastatic Breast Cancer Project (www.mbcproject.org) (n=18) and the Metastatic Melanoma (n=38)^84^ databases were downloaded from cBioPortal (www.cbioportal.org) on February 8, 2023. Average log_2_ expression of the mTORC1^ECKO^ and cytotoxic T lymphocyte (CTL) gene sets (Extended Data Table 4) comprise each gene signature, respectively.

Single-cell RNA-seq data from breast cancer (GSE176078)^85^, lung adenocarcinoma (Bischoff et al.)^86^, and melanoma (GSE72056)^87^ patient datasets were used to analyze previously identified cell populations with Seurat v.4.9.9^88^. Gene set enrichment scores for endothelial cells were determined by performing ssGSEA on each single-cell expression matrix using GSVA in R (v.4.3.1)^89^. REACTOME gene sets for “mTORC1-mediated signaling” and “RAB regulation of trafficking” (Extended Data Table 5) were identified from the curated C2 collection of the Molecular Signatures Database (MSigDB)^81, 90, 91^. Individual endothelial cells were stratified based on the median mTORC1-mediated signaling enrichment score and *CLSTN1* expression was determined.

### Survival analysis

Recurrence-free survival (RFS), first progression survival (FP), or progression-free survival (PFS) data in breast cancer^92^, lung cancer^93^, and melanoma^94^, respectively, was downloaded from KM plot (www.kmplot.com). Patients were stratified by mean expression of genes comprising the mTORC1^ECKO^ gene set in Extended Data Table 4, using the following expression cutoffs: breast cancer RFS (674.7, range=265-2705), lung cancer FP (1132.65, range=312-2733), no ICI melanoma PFS (1060.1, range=488-5000), and Pembrolizumab melanoma PFS (1165.49, range=557-2805).

### Statistics

All plots and statistical analyses were performed using GraphPad Prism software (10.0.3). Individual data points are shown where possible. Summary data are mean with SEM. For in vivo tumor experiments, data are reported from 2-4 independent experiments, where each point represents individual animals. Statistical comparisons between two groups were performed using unpaired Student’s t-test or Welch’s t-test, as indicated. For multiple comparisons, one- or two-way analysis of variance (ANOVA) was performed with individual comparisons evaluated using Tukey’s, Dunnett’s, or Sidak’s post hoc analysis, as indicated. Outliers were excluded using the ROUT method (Q=5%). All statistical analyses were two-tailed, and differences with a p-value less than 0.05 were considered to be statistically significant.

For Kaplan-Meier survival curve analysis, log-rank analysis was performed between groups using the Mantel-Cox method. Hazard ratios (HR) and 95% confidence intervals (CI) were determined using the log-rank test. Pearson correlation (r) was performed on TCGA datasets.

### Study approval

Studies involving animals were performed with approval from the Vanderbilt University Medical Center’s Institutional Animal Care and Utilization Committee (IACUC).

## Acknowledgements

We thank Drs. Ralf Adam (Max Planck Institute) and Dr. Hong Chen (Boston Children’s Hospital, Harvard Medical School) for providing the CDH5-Cre^ER^ transgenic mice used in this study, and Dr. Brad Reinfeld for thoughtful discussion. This work is supported by a NIH R01 CA271176, NIH R01 CA095004, and VA Career Scientist Award 5IK6BX005391 to J.C. J.C. is also supported by NIH R01 CA250506 and VA Merit Award 5101BX000134. D.N.E. is supported by a Department of Defense CDMRP W81XWH2210109. L.C.K. was supported by NIH T32 CA009592 and NIH F31 CA220804 and is currently supported by NCI T32 CA09140 and a postdoctoral fellowship from the American Cancer Society (PF-23-1034739-01-TBE). V.M.N. is supported by NIH T32 CA009592. K.C.E. is supported by NIH 1R011NS118580. Flow cytometry experiments were performed in the Vanderbilt University Medical Center (VUMC) Flow Cytometry Shared Resource, which is supported by the Vanderbilt Ingram Cancer Center (P30 CA68485) and the Vanderbilt Digestive Disease Research Center (DK058404). Confocal imaging was performed in part through the use of the Vanderbilt Cell Imaging Shared Resource (supported by NIH grants CA68485, DK20593, DK58404, DK59637, and EY08126). Tissue processing and H&E staining was performed by the VUMC Translational Pathology Shared Resource, which is supported by NCI/NIH Cancer Center Support Grant P30CA068485.

## Contributions

D.N.E., S.W., and J.C. conceptualized the project and developed methodologies. D.N.E., S.W., W.S., L.C.K., V.M.N., and Y.H. performed experiments. Data analysis and interpretation was performed by D.N.E., S.W., and J.C. K.C.E. and M.R.B. provided critical feedback and reagents. D.N.E. and J.C. wrote the manuscript. D.N.E., S.W., W.S., L.C.K., K.C.E., M.R.B., and J.C. reviewed and/or revised the manuscript.

## Data Availability

Mouse RNA-sequencing data generated by this study has been deposited in the Gene Expression Omnibus (GEO) and are available under the accession numbers GSE256508 (tumor cells) and GSE256509 (endothelial cells). Publicly available single-cell RNA sequencing data used in this study can be found under the accession numbers GSE176078 and GSE72056, or at Code Ocean (doi: 10.24433/CO.0121060.v1). Code used in this study for analysis of single-cell RNA sequencing databases is available at https://github.com/dne7754/EC_sc_RAB_Cal1/tree/main. Source data for in vitro and in vivo experiments in Figures 1-5 and Extended Data Figures, metabolic data, and full western blot source images are included in Source Data.

