## Supplementary Data for "Regulation of fatty acid delivery to metastases by tumor endothelium"

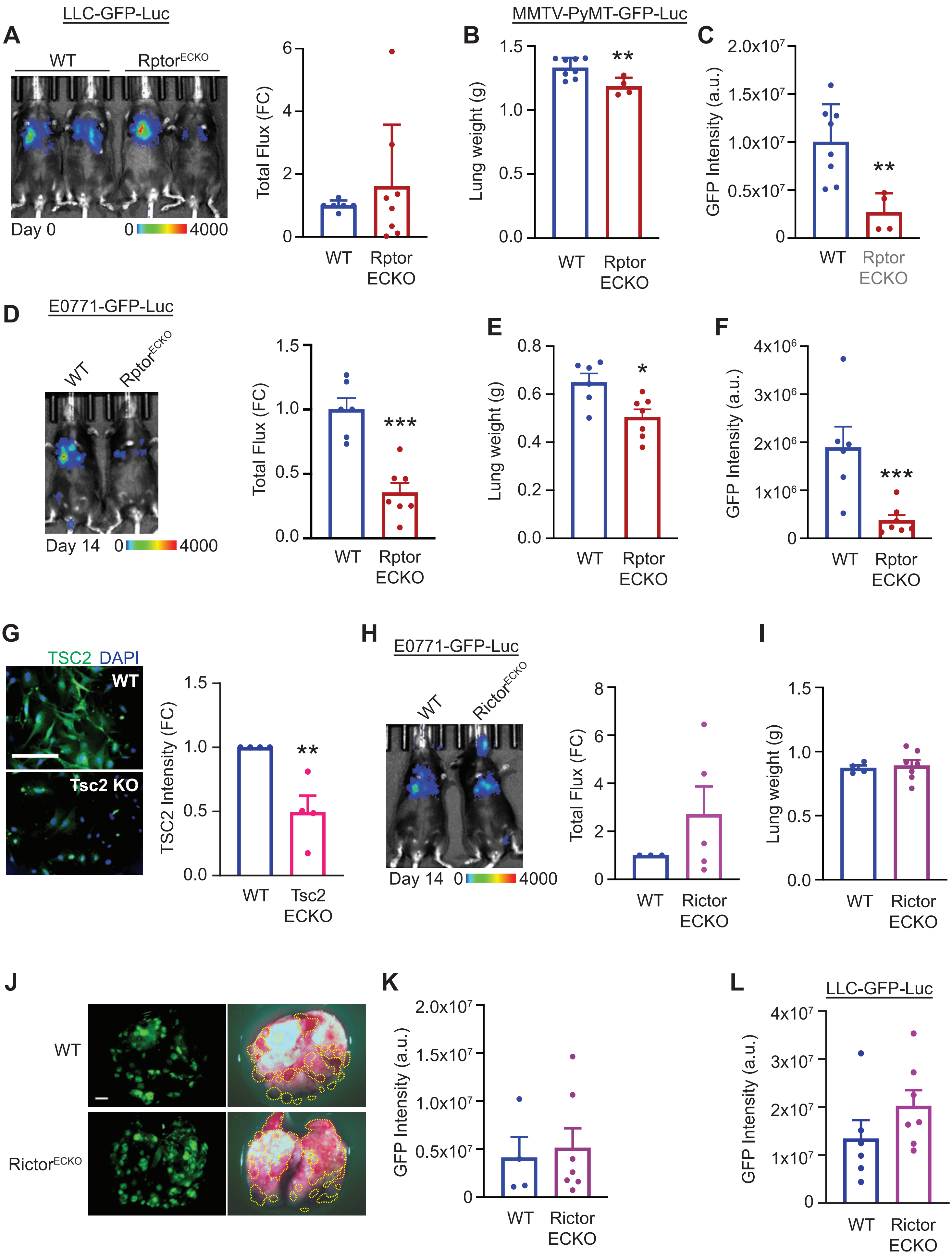

LLC-GFP-Luc
A
WT
Rptor^ECKO
Day 0
0
4000
Total Flux (FC)
6
4
2
0
WT
Rptor ECKO
MMTV-PyMT-GFP-Luc
B
Lung weight (g)
1.5
1.0
0.5
0.0
**
WT
Rptor ECKO
C
GFP Intensity (a.u.)
2.0x10^7
1.5x10^7
1.0x10^7
0.5x10^7
0
**
WT
Rptor ECKO
D
E0771-GFP-Luc
WT
Rptor^ECKO
Day 14
0
4000
Total Flux (FC)
1.5
1.0
0.5
0.0
***
WT
Rptor ECKO
E
Lung weight (g)
0.8
0.6
0.4
0.2
0.0
*
WT
Rptor ECKO
F
GFP Intensity (a.u.)
4x10^6
3x10^6
2x10^6
1x10^6
0
***
WT
Rptor ECKO
G
TSC2 DAPI
WT
Tsc2 KO
TSC2 Intensity (FC)
1.5
1.0
0.5
0.0
**
WT
Tsc2 ECKO
H
E0771-GFP-Luc
WT
Rictor^ECKO
Day 14
0
4000
Total Flux (FC)
8
6
4
2
0
WT
Rictor ECKO
I
Lung weight (g)
1.5
1.0
0.5
0.0
WT
Rictor ECKO
J
WT
Rictor^ECKO
K
GFP Intensity (a.u.)
2.0x10^7
1.5x10^7
1.0x10^7
0.5x10^7
0
WT
Rictor ECKO
L
LLC-GFP-Luc
GFP Intensity (a.u.)
4x10^7
3x10^7
2x10^7
1x10^7
0
WT
Rictor ECKO

**Extended Data Figure 1. Loss of Raptor/mTORC1, but not Rictor/mTORC2, in endothelium reduces metastatic outgrowth of cancer cells in the lung. (A)** Representative bioluminescence images from Day 0 of LLC-GFP-luc inoculated WT or $Rptor^{ECKO}$ male mice. Scale bar shows counts. Total radiance flux was normalized to WT controls. Unpaired t-test, p=0.472. **(B-D)** WT or $Rptor^{ECKO}$ female mice were inoculated with MMTV-PyMT-GFP-luc cells, followed by tamoxifen treatment starting on day 4. Tumors were harvested on day 14. (**B)** Lung weights were recorded in grams (g) at harvest and **(C)** GFP intensity was calculated as arbitrary units (a.u.). Unpaired t-test, $p=9.75x10^{-3}$ for lung weights and $p=6.30x10^{-3}$ for GFP intensity. **(D-F)** WT or $Rptor^{ECKO}$ female mice were inoculated with E0771-GFP-luc cells as described in (B). **(D)** Representative bioluminescence images are shown from day 14. Scale bar shows counts. Total radiance flux was normalized to WT controls. Unpaired t-test, $p=1.65x10^{-3}$. Tumors were harvested on day 18. **(E)** Lung weights were recorded in grams (g) at harvest and **(F)** GFP intensity was calculated. Unpaired t-test, p=0.0131 for lung weights and $p=3.97x10^{-3}$ for GFP intensity. **(G)** Primary microvascular endothelial cells were isolated from $Tsc2^{fl/fl}$ mice were transduced with control (WT) or cre-expressing (Tsc2 KO) adenovirus. Immunofluorescence was used to confirm Tsc2 deletion. Representative images of TSC2 (green) are shown. Nuclei are stained with DAPI (blue). Scale bar is 100 $\mu$m. Fluorescence intensity was normalized to WT control. Unpaired t-test, $p=8.11x10^{-3}$. **(H-K)** WT or $Rictor^{ECKO}$ female mice were inoculated with E0771-GFP-luc cells as described above. **(H)** Representative bioluminescence images from day 14 are shown. Scale bar shows counts. Total radiance flux was normalized to WT controls. Unpaired t-test, p=0.318. **(I)** Tumors were harvested on day 18, and lung weights were recorded in grams (g). Unpaired t-test, p=.767. **(J)** representative GFP (left) and gross (right) lung images are shown. Scale bar is 5 mm. Visible tumor area is outlined by yellow line. **(K)** GFP intensity was calculated as arbitrary units (a.u.). Unpaired t-test, p=0.753. **(L)** WT or $Rictor^{ECKO}$ male mice were inoculated with LLC-GFP-luc cells as described above. Tumors were harvested on day 18. Lung weights were recorded. Unpaired t-test, p=0.204. *p<0.05, **p<0.01, ***p<0.005.

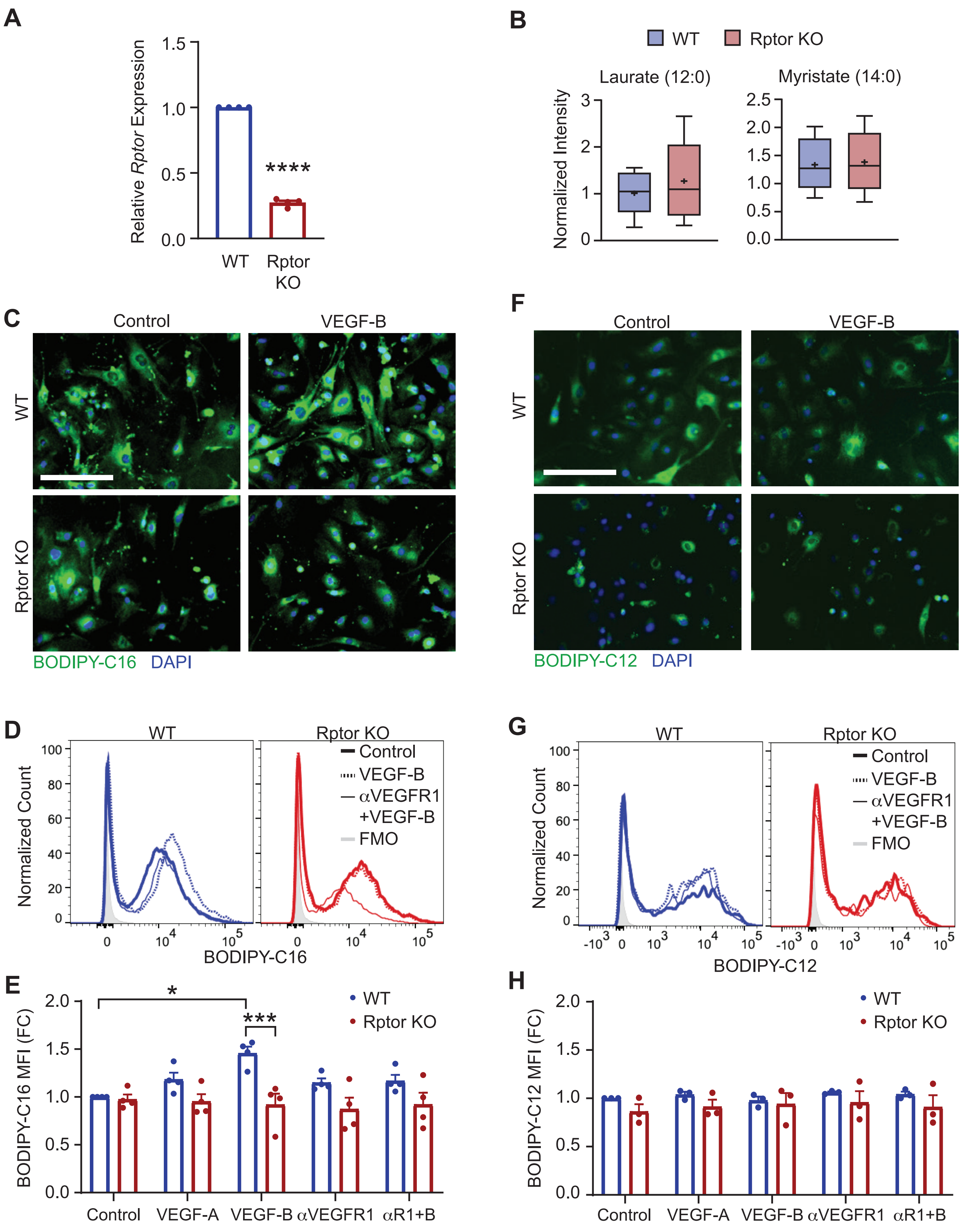

A
Relative Rptor Expression
1.5
1.0
0.5
0.0
****
WT
Rptor KO
B
WT
Rptor KO
Laurate (12:0)
Myristate (14:0)
Normalized Intensity
C
Control
VEGF-B
WT
Rptor KO
BODIPY-C16 DAPI
F
Control
VEGF-B
WT
Rptor KO
BODIPY-C12 DAPI
D
WT
Rptor KO
Normalized Count
Control
VEGF-B
αVEGFR1
+VEGF-B
FMO
BODIPY-C16
G
WT
Rptor KO
Normalized Count
Control
VEGF-B
αVEGFR1
+VEGF-B
FMO
BODIPY-C12
E
BODIPY-C16 MFI (FC)
*
***
WT
Rptor KO
Control
VEGF-A
VEGF-B
αVEGFR1
αR1+B
H
BODIPY-C12 MFI (FC)
WT
Rptor KO
Control
VEGF-A
VEGF-B
αVEGFR1
αR1+B

**Extended Data Figure 2. Targeting Raptor/mTORC1 reduces fatty acid long-chain fatty acid content in endothelial cells. (A-B)** Metabolomics was performed on primary microvascular endothelial cells isolated from Rptor$^{fl/fl}$ mice transduced with control (WT) or cre recombinase (KO) adenoviruses. **(A)** Rptor deletion was confirmed by reduced Rptor RNA expression by qRT-PCR. Unpaired t-test, $p=6.19x10^{-9}$. **(B)** Normalized intensities ($log_2$+1) of representative medium chain fatty acid (MCFA) metabolites. Welch's t-test; 12:0, p=0.533; 14:0, p=0.879. **(C-H)** Primary microvascular endothelial cells were isolated from Rptor$^{fl/fl}$ mice, and deletion was carried out by transduction with control (WT) or cre-expressing (KO) adenovirus. **(C)** Cells were treated without (Control) or with VEGF-B (300 ng/mL) for 30 hours in free fatty acid BSA (FFA-BSA) supplemented media, followed by incubation with BODIPY FL C16 (BODIPY-C16, 20 μg/mL) for 3 min. Representative images of BODIPY-C16 (green) are shown. Nuclei were stained with DAPI (blue). Scale bar is 100 μm. **(D-E)** Cells were treated with VEGF-B (300 ng/mL), VEGF-A (300 ng/mL), anti-VEGFR1 (1 μg/mL), or anti-VEGFR1+VEGF-B (αR1+B) and analyzed by flow cytometry. **(D)** Representative histogram plots of Control, VEGF-B, and anti-VEGFR1+VEGF-B groups are shown. **(E)** BODIPY-C16 median fluorescence intensity (MFI) was normalized to WT Control. Two-way ANOVA ($p=1.56x10^{-5}$) with Tukey's post hoc. **(F-H)** Cells were treated with BODIPY-C12 (20 μg/mL) as described in (A-C). **(F)** Representative images of BODIPY-C12 (green) are shown. Nuclei were stained with DAPI (blue). Scale bar is 100 μm. **(G)** Representative histogram plots of Control, VEGF-B, and anti-VEGFR1+VEGF-B are shown. **(H)** BODIPY-C12 MFI was normalized to WT Control. Two-way ANOVA (p=0.0339) with Tukey's post hoc, which revealed no significant comparisons. *p<0.05, ***p<0.005, ****p<0.001.

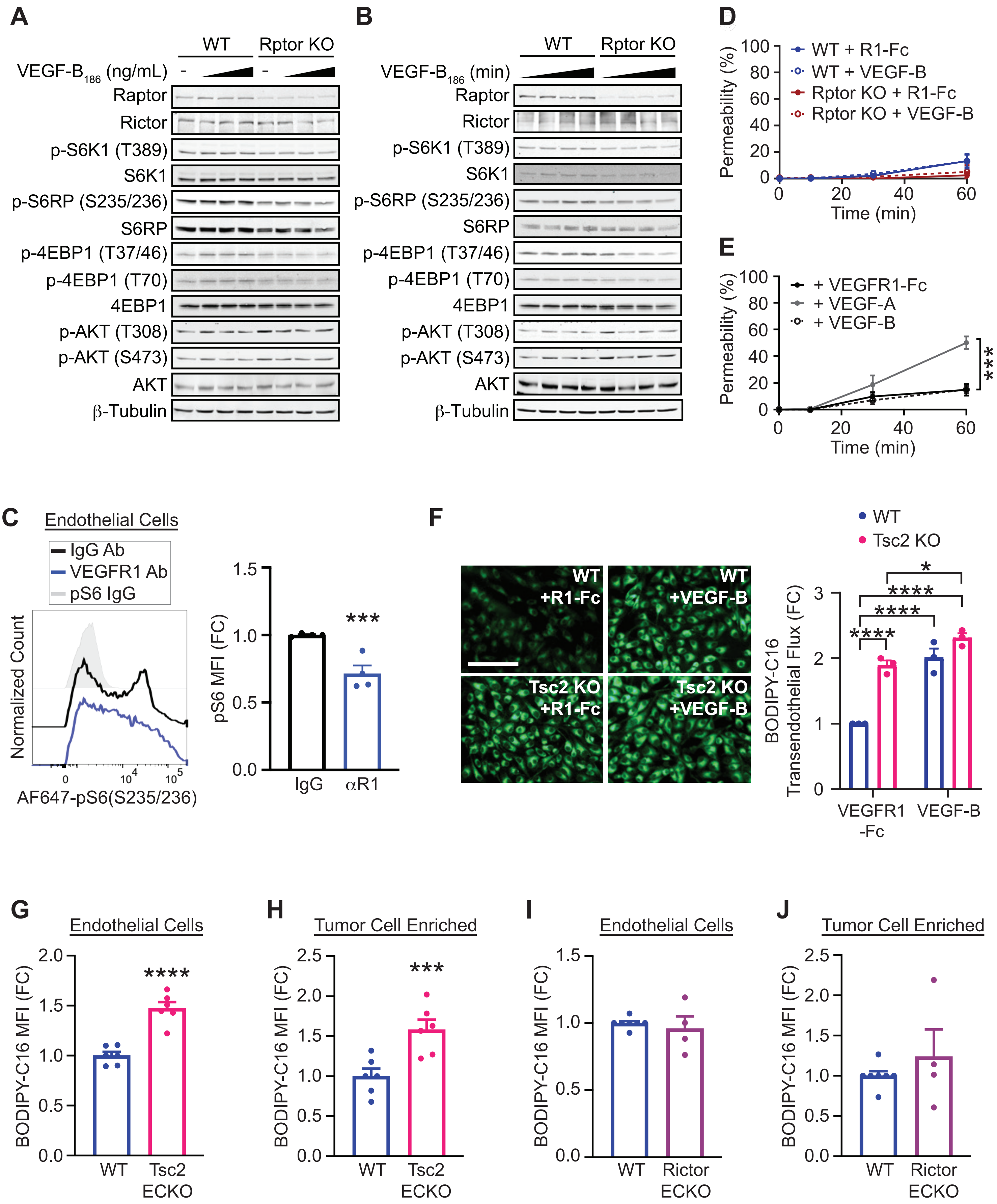

A
WT
Rptor KO
VEGF-B186 (ng/mL)
Raptor
Rictor
p-S6K1 (T389)
S6K1
p-S6RP (S235/236)
S6RP
p-4EBP1 (T37/46)
p-4EBP1 (T70)
4EBP1
p-AKT (T308)
p-AKT (S473)
AKT
β-Tubulin
B
WT
Rptor KO
VEGF-B186 (min)
D
WT + R1-Fc
WT + VEGF-B
Rptor KO + R1-Fc
Rptor KO + VEGF-B
Permeability (%)
Time (min)
E
+ VEGFR1-Fc
+ VEGF-A
+ VEGF-B
***
C
Endothelial Cells
IgG Ab
VEGFR1 Ab
pS6 IgG
Normalized Count
AF647-pS6(S235/236)
pS6 MFI (FC)
IgG
αR1
***
F
WT +R1-Fc
WT +VEGF-B
Tsc2 KO +R1-Fc
Tsc2 KO +VEGF-B
WT
Tsc2 KO
BODIPY-C16 Transendothelial Flux (FC)
VEGFR1 -Fc
VEGF-B
G
Endothelial Cells
BODIPY-C16 MFI (FC)
WT
Tsc2 ECKO
****
H
Tumor Cell Enriched
***
I
Endothelial Cells
WT
Rictor ECKO
J
Tumor Cell Enriched

**Extended Data Figure 3. Endothelial Raptor/mTORC1 promotes transendothelial transport of fatty acids in response to VEGF-B. (A-B)** Primary microvascular endothelial cells were isolated from Rptor$^{fl/fl}$ mice, and deletion was carried out by transduction with control (WT) or cre-expressing (KO) adenovirus. **(A)** Cells were serum starved overnight and stimulated with VEGF-B (0, 75, 150, or 300 ng/mL) for 30 hours. Lysates were harvested for signaling analysis by immunoblotting. **(B)** Cells were serum starved as described in (A) and treated with VEGF-B (300 ng/mL) for 0, 5, 15, or 30 min. **(C)** Mice bearing E0771-GFP-luc lung metastases were treated with anti-VEGFR1 antibody or IgG control. CD31+ endothelial cells from harvested tumors were analyzed for pS6 (S235/236) by flow cytometry. Unpaired t-test, p= 0.003259. **(D-E)** Endothelial permeability assay was performed by detecting diffusion of Texas Red (TR)-Dextran across a transwell coated with a confluent top layer of WT or Rptor KO primary microvascular endothelial cells. Media was removed from the upper chamber at 0, 10, 30, and 60 min and fluorescence was analyzed on a plate reader. **(F)** Transendothelial transport assay of WT or Tsc2 KO primary vascular endothelial cells, as described in Figure 3. Representative images of BODIPY-C16 (green) in LLC tumor cells. Scale bar is 100 μm. BODIPY-C16 intensity was normalized to WT + VEGFR1-Fc control. Two-way ANOVA ($p=8.04x10^{-3}$) with Tukey's post hoc. **(G-H)** WT or Tsc2$^{ECKO}$ female mice were inoculated with E0771-luc cells and injected with BODIPY FL C16 (BODIPY-C16), as described in Figure 3. BODIPY-C16 median fluorescence intensity (MFI) was determined by flow cytometry in (G) $CD45^-CD31^+$ endothelial cells and (H) $CD45^-CD31^-FSC^{hi}$ tumor-cell enriched populations and normalized to WT controls. Representative histograms are shown. Unpaired t-test; $CD31^+$, $p=6.26x10^{-5}$; tumor-cell enriched, $p=3.90x10^{-3}$. **(I-J)** WT or Rictor$^{ECKO}$ male mice were inoculated with LLC cells and injected with BODIPY FL C16 (BODIPY-C16), as described above. BODIPY-C16 median fluorescence intensity (MFI) was determined by flow cytometry in (I) $CD45^-CD31^+$ endothelial cells and (J) $CD45^-CD31^-FSC^{hi}$ tumor-cell enriched populations and normalized to WT controls. Representative histograms are shown. Unpaired t-test; $CD31^+$, p=0.577; tumor-cell enriched, p=0.376. *p<0.05, **p<0.01, ***p<0.005, ****p<0.001.

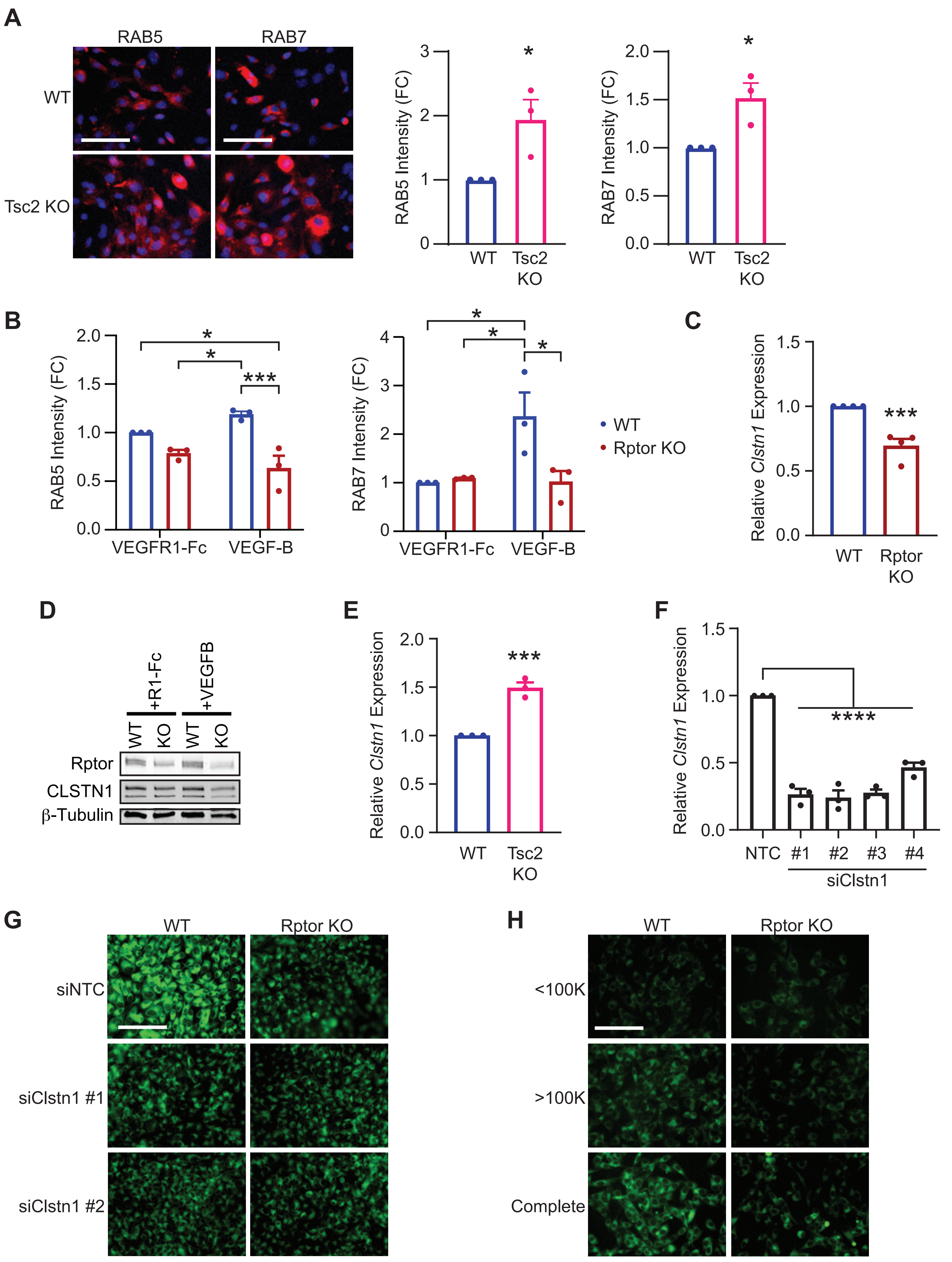

A
RAB5
RAB7
WT
Tsc2 KO
RAB5 Intensity (FC)
RAB7 Intensity (FC)
WT
Tsc2 KO
*
B
RAB5 Intensity (FC)
RAB7 Intensity (FC)
VEGFR1-Fc
VEGF-B
*
***
WT
Rptor KO
C
Relative Clstn1 Expression
WT
Rptor KO
***
D
+R1-Fc
+VEGFB
WT
KO
Rptor
CLSTN1
β-Tubulin
E
Relative Clstn1 Expression
WT
Tsc2 KO
***
F
Relative Clstn1 Expression
NTC
#1
#2
#3
#4
siClstn1
****
G
WT
Rptor KO
siNTC
siClstn1 #1
siClstn1 #2
H
WT
Rptor KO
<100K
>100K
Complete

**Extended Data Figure 4. mTORC1 increases RAB endosomes and regulates CLSTN1 expression to support transendothelial transport of fatty acids in endothelial cells. (A)** Immunofluorescence of RAB5 (left) or RAB7 (right) was performed on WT or TSC2 KO primary microvascular endothelial cells cultured in complete endothelial cell media. Representative immunofluorescence images of RAB5 and RAB 7 are shown (both red). Nuclei are stained with DAPI (blue). Scale bars are 100 μm. RAB5 or RAB7 intensities were normalized to WT controls. Unpaired t-test; RAB5, p=0.0371; RAB7, p=0.0255. **(B)** WT or Rptor KO primary microvascular endothelial cells cultured in basal media supplemented with FFA-BSA were stimulated with VEGFR1-Fc (1 μg/mL) or VEGF-B (300 ng/mL) for 30 hours. Immunofluorescence and intensities were performed as in (A). Two-way ANOVA (RAB5, p=0.0372; RAB7, p=0.0283) with Tukey's post hoc. **(C-D)** WT or Rptor KO primary microvascular endothelial cells were assessed for *Clstn1* expression by (C) qRT-PCR and (D) immunoblotting. (C) Unpaired t-test, $p=1.27x10^{-3}$. **(E)** *Clstn1* expression was analyzed in WT or Tsc2 KO primary microvascular endothelial cells by qRT-PCR. Unpaired t-test, $p=9.96x10^{-4}$. **(F)** Knockdown of *Clstn1* using siRNA was confirmed by reduced *Clstn1* expression in $Rptor^{fl/fl}$ primary microvascular endothelial cells treated with control adenovirus. One-way ANOVA ($p=1.84x10^{-7}$) with Dunnett's post hoc. **(G)** Representative images of LLC tumor cells from transendothelial transport of BODIPY-C16 (green) in non-targeting control (NTC) or CLSTN1 knockdown endothelial cells from Figure 4. Scale bar is 100 μm. **(H)** Representative images of LLC tumor cells cultured in small (<100 kDa) or large (>100 kDa) fractionated endothelial cell conditioned media from Figure 4. Images from complete conditioned media are also shown. Scale bar is 100 μm. *p<0.05, ***p<0.005, ****p<0.001.

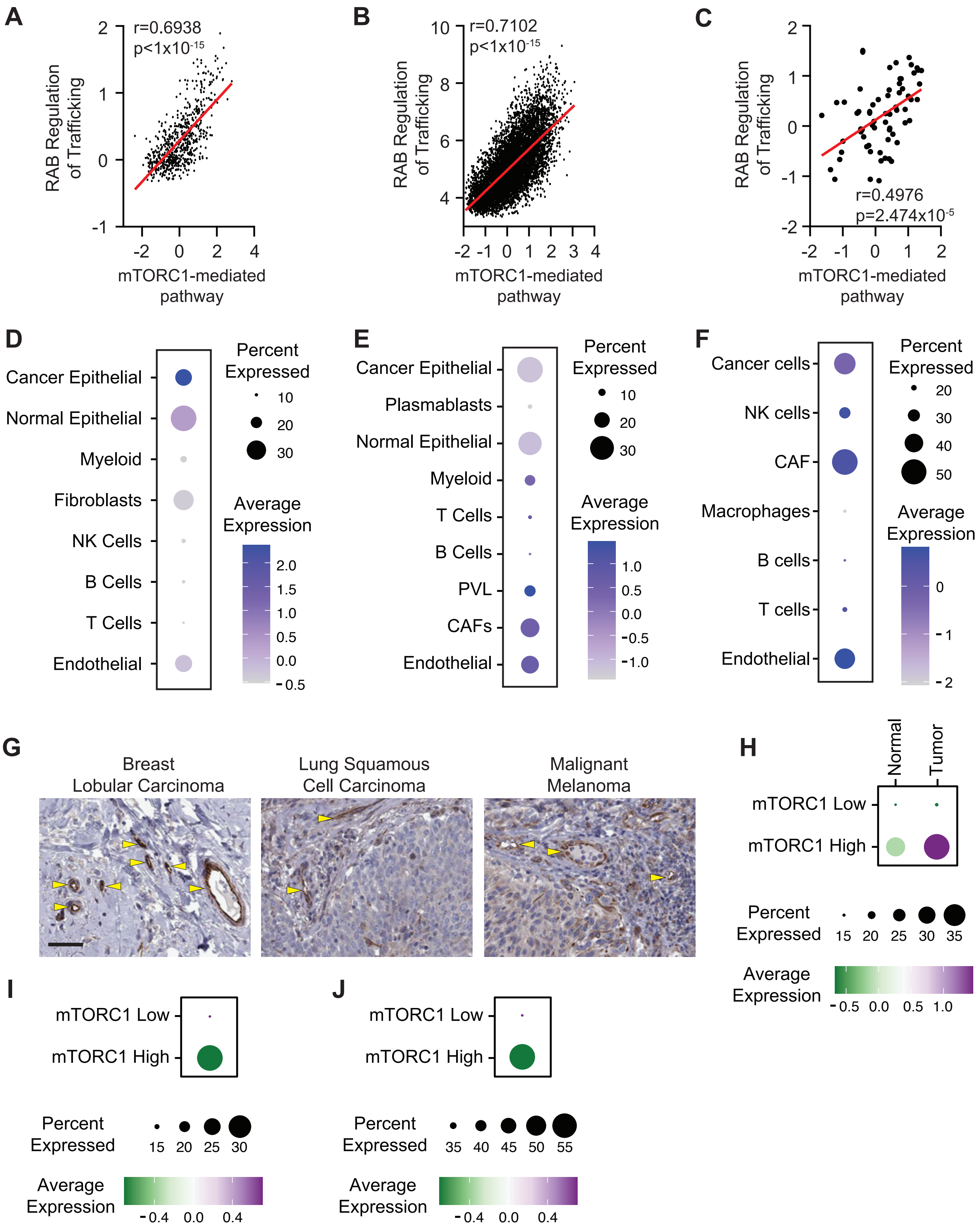

A
r=0.6938
p<1x10-15
RAB Regulation of Trafficking
mTORC1-mediated pathway
B
r=0.7102
p<1x10-15
RAB Regulation of Trafficking
mTORC1-mediated pathway
C
r=0.4976
p=2.474x10-5
RAB Regulation of Trafficking
mTORC1-mediated pathway
D
Cancer Epithelial
Normal Epithelial
Myeloid
Fibroblasts
NK Cells
B Cells
T Cells
Endothelial
Percent Expressed
Average Expression
E
Cancer Epithelial
Plasmablasts
Normal Epithelial
Myeloid
T Cells
B Cells
PVL
CAFs
Endothelial
Percent Expressed
Average Expression
F
Cancer cells
NK cells
CAF
Macrophages
B cells
T cells
Endothelial
Percent Expressed
Average Expression
G
Breast Lobular Carcinoma
Lung Squamous Cell Carcinoma
Malignant Melanoma
H
Normal
Tumor
mTORC1 Low
mTORC1 High
Percent Expressed
Average Expression
I
mTORC1 Low
mTORC1 High
Percent Expressed
Average Expression
J
mTORC1 Low
mTORC1 High
Percent Expressed
Average Expression

**Extended Data Figure 5. mTORC1 signaling correlates with RAB trafficking and *CLSTN1* expression in human tumor-associated endothelial cells. (A-C)** mTORC1 signaling positively correlates with RAB regulation of trafficking gene set in endothelial cells from the (A) lung adenocarcinoma (Bischoff, et al.), (B) breast cancer (GSE176078), and (C) melanoma (GSE72056) single-cell RNA-seq datasets. The x and y axis represents z-scores of ssGSEA enrichment of indicated gene sets. **(D-E)** Dot plots of *CLSTN1* expression in cell populations from (D) lung adenocarcinoma, (E) breast cancer, and (F) melanoma datasets above. **(G)** Immunohistochemistry of CLSTN1 from patient samples showing strong staining in vascular endothelial regions, denoted by yellow arrows. Images were acquired from the Human Protein Atlas. **(H)** Dot plot of *CLSTN1* expression in normal or tumor-associated endothelial cells, stratified based on mTORC1 signaling ssGSEA enrichrment scores, in the lung adenocarcinoma dataset used above. **(I-J)** Dot plot of *CLSTN1* expression in tumor-associated endothelial cells in (I) breast cancer and (J) melanoma datasets described above. mTORC1 signaling stratification was performed as described in (H).

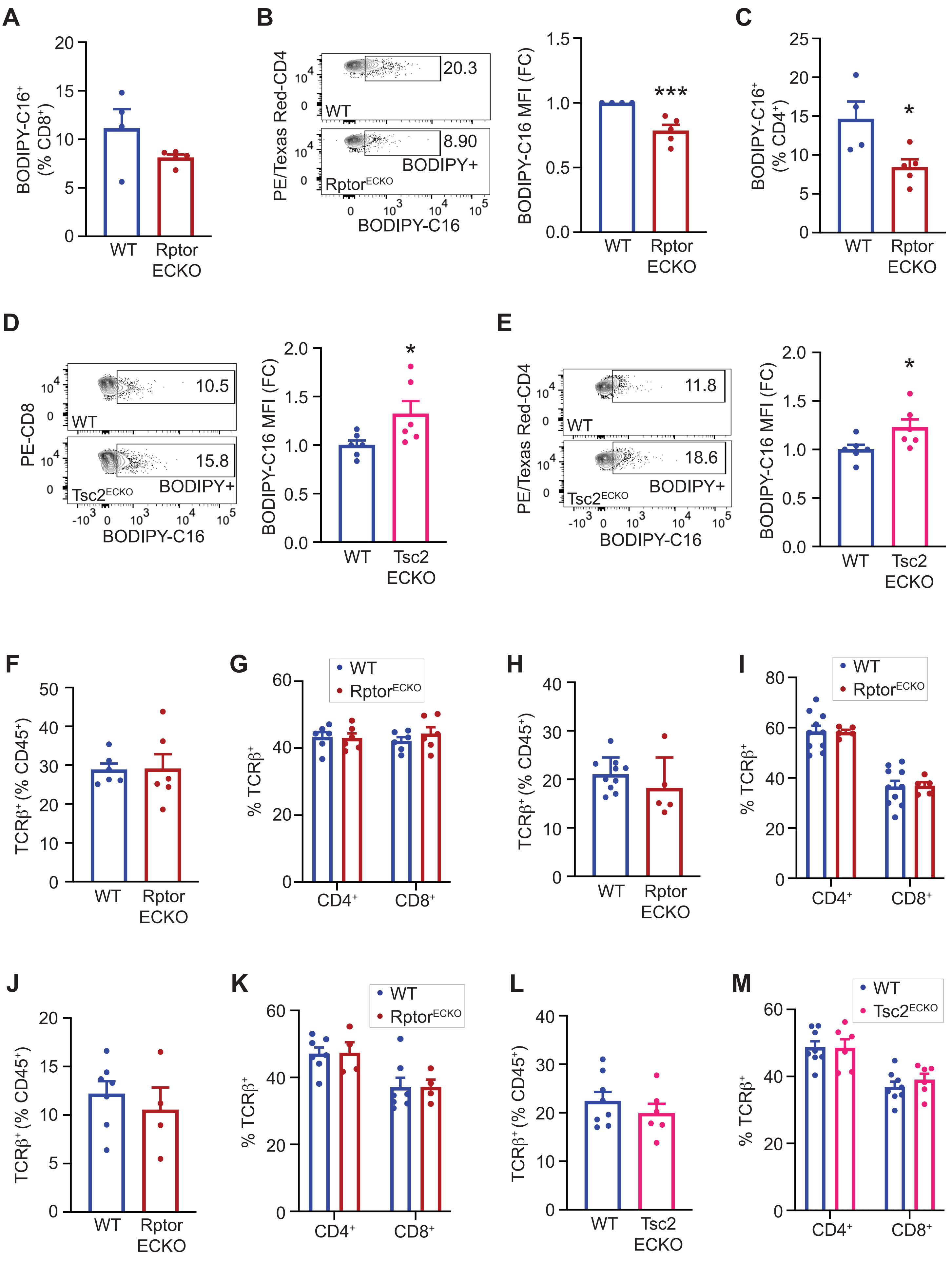
A
B
C
D
E
F
G
H
I
J
K
L
M
BODIPY-C16⁺ (% CD8⁺)
BODIPY-C16⁺ (% CD4⁺)
PE/Texas Red-CD4
PE-CD8
BODIPY-C16
BODIPY+
BODIPY-C16 MFI (FC)
20.3
8.90
10.5
15.8
11.8
18.6
WT
Rptor ECKO
Rptor^ECKO
Tsc2 ECKO
Tsc2^ECKO
TCRβ⁺ (% CD45⁺)
% TCRβ⁺
CD4⁺
CD8⁺
***
*

**Extended Data Figure 6. Endothelial mTORC1 promotes BODIPY-C16 uptake in tumor-associated T cells. (A-C)** WT or $Rptor^{ECKO}$ male mice were inoculated with LLC tumors cells and injected with BODIPY-C16, as described in Figure 5. BODIPY-C16+ cells are shown as (A) a percent of $CD8^+$ T cells, (B) median fluorescence intensity (MFI) in $CD4^+$ T cells, or (C) percent of $CD4^+$ T cells. Representative plots of BODIPY-C16 in $CD4^+$ T cells are shown. BODIPY-C16 MFI was normalized to WT controls, presented as fold change (FC). Unpaired t-test, BODIPY+ (% $CD8^+$), p=0.132; $CD4^+$ MFI, $p=4.33 \times 10^{-3}$; BODIPY+ (% $CD4^+$), p=0.030. **(D-E)** WT or $TSC2^{ECKO}$ female mice were inoculated with E0771-luc tumor cells and injected with BODIPY-C16, as in (A-C). Representative plots of BODIPY-C16 in (D) $CD8^+$ T cells or (E) $CD4^+$ T cells are shown. BODIPY-C16 median fluorescence intensity (MFI) was calculated and normalized to WT controls, presented as fold change (FC). Unpaired t-test; $CD8^+$ T cells, p=0.0464; $CD4^+$ T cells, p=0.0422. **(F-G)** WT or $Rptor^{ECKO}$ female mice were inoculated with E0771-GFP-luc tumor cells. Metastatic tumors were analyzed by flow cytometry. (F) T cells ($TCR\beta^+$) and (G) $CD4^+$/$CD8^+$ T cells are shown as percentage (%) of $CD45^+$ or $TCR\beta^+$, respectively. Unpaired t-test: $TCR\beta^+$, p=0.946; $CD4^+$, p=0.868; $CD8^+$, p=0.369. **(H-I)** WT or $Rptor^{ECKO}$ female mice were inoculated with MMTV-PyMT-GFP-luc tumor cells. Metastatic tumors were analyzed for (H) T cells ($TCR\beta^+$) and (I) $CD4^+$/$CD8^+$ T cells, as described in (F-G). Unpaired t-test: $TCR\beta^+$, p=0.275; $CD4^+$, p=0.960; $CD8^+$, p=0.932. **(J-K)** WT or $Rptor^{ECKO}$ male mice were inoculated with LLC-GFP-luc tumor cells. Metastatic tumors were analyzed for (J) T cells ($TCR\beta^+$) and (K) $CD4^+$/$CD8^+$ T cells, as described above. Unpaired t-test: $TCR\beta^+$, p=0.515; $CD4^+$, p=0.944; $CD8^+$, p=0.997. **(L-M)** WT or $Tsc2^{ECKO}$ female mice were inoculated with E0771-GFP-luc tumor cells. Metastatic tumors were analyzed for (L) T cells ($TCR\beta^+$) and (M) $CD4^+$/$CD8^+$ T cells, as described above. Unpaired t-test: $TCR\beta^+$, p=0.376; $CD4^+$, p=0.958; $CD8^+$, p=0.396. *p<0.05, ***p<0.005.

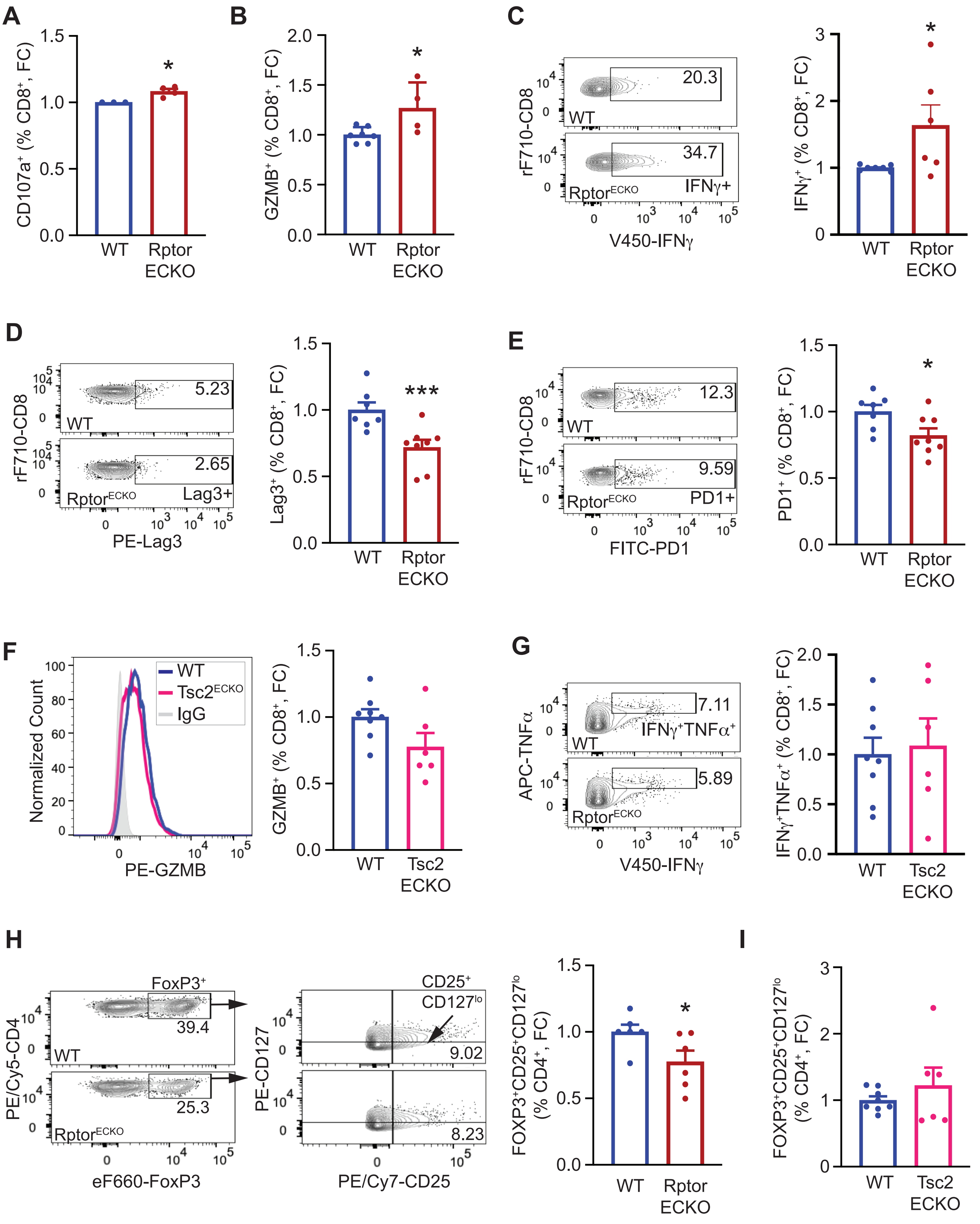

A
CD107a+ (% CD8+, FC)
WT
Rptor ECKO
*
B
GZMB+ (% CD8+, FC)
WT
Rptor ECKO
*
C
rF710-CD8
V450-IFNγ
20.3
34.7
WT
RptorECKO
IFNγ+
IFNγ+ (% CD8+, FC)
*
D
rF710-CD8
PE-Lag3
5.23
2.65
WT
RptorECKO
Lag3+
Lag3+ (% CD8+, FC)
***
E
rF710-CD8
FITC-PD1
12.3
9.59
WT
RptorECKO
PD1+
PD1+ (% CD8+, FC)
*
F
Normalized Count
PE-GZMB
WT
Tsc2ECKO
IgG
GZMB+ (% CD8+, FC)
WT
Tsc2 ECKO
G
APC-TNFα
V450-IFNγ
7.11
5.89
IFNγ+TNFα+
WT
RptorECKO
IFNγ+TNFα+ (% CD8+, FC)
WT
Tsc2 ECKO
H
PE/Cy5-CD4
eF660-FoxP3
FoxP3+
39.4
25.3
WT
RptorECKO
PE-CD127
PE/Cy7-CD25
CD25+ CD127lo
9.02
8.23
FOXP3+CD25+CD127lo (% CD4+, FC)
WT
Rptor ECKO
*
I
FOXP3+CD25+CD127lo (% CD4+, FC)
WT
Tsc2 ECKO

**Extended Data Figure 7. Endothelial Raptor/mTORC1 loss improves anti-tumor immune activity of $CD8^+$ T cells. (A)** WT or $Rptor^{ECKO}$ female mice were inoculated with E0771 cells and lung metastatic tumors were analyzed by flow cytometry, as described in Figure 5. $CD8^+CD107^+$ cells were normalized to WT controls and shown as fold change (FC). Unpaired t-test, $p=0.0144$. **(B)** WT or $Rptor^{ECKO}$ female mice were inoculated with MMTV-PyMT cells and lung metastatic tumors were analyzed by flow cytometry. $CD8^+GZMB^+$ cells were normalized, as described in (A). Unpaired t-test, $p=0.0274$. **(C)** WT or $Rptor^{ECKO}$ male mice were inoculated with LLC tumor cells and lung metastatic tumors were analyzed by flow cytometry. $CD8^+IFN\gamma^+$ cells were normalized as in (A). Unpaired t-test, $p=0.0359$. **(D-E)** Metastatic tumor samples from (A) were analyzed for (D) Lag3 or (E) PD1. Unpaired t-test; $Lag3^+$, $p=0.00415$; $PD1^+$, $p=0.0303$. **(F-G)** WT or $Tsc2^{ECKO}$ female mice were inoculated and analyzed for (F) GZMB or (G) $IFN\gamma/TNF\alpha$, as described in (A). Unpaired t-test; $GZMB^+$, $p=0.0709$; $IFN\gamma^+TNF\alpha^+$, $p=0.781$. **(H)** WT or $Rptor^{ECKO}$ female mice were inoculated with E0771 tumor cells, and lung metastatic tumors were analyzed by flow cytometry, as described in Figure 5. $CD4^+FoxP3^+$ Tregs were assessed for the activation markers $CD25^+CD127^{lo}$. Activated Tregs were normalized to WT controls and shown as fold change (FC). Unpaired t-test, $p=0.0443$. **(I)** WT or $Tsc2^{ECKO}$ female mice were inoculated and analyzed as in (H). Unpaired t-test, $p=0.375$. $^*p<0.05$, $^{***}p<0.005$.

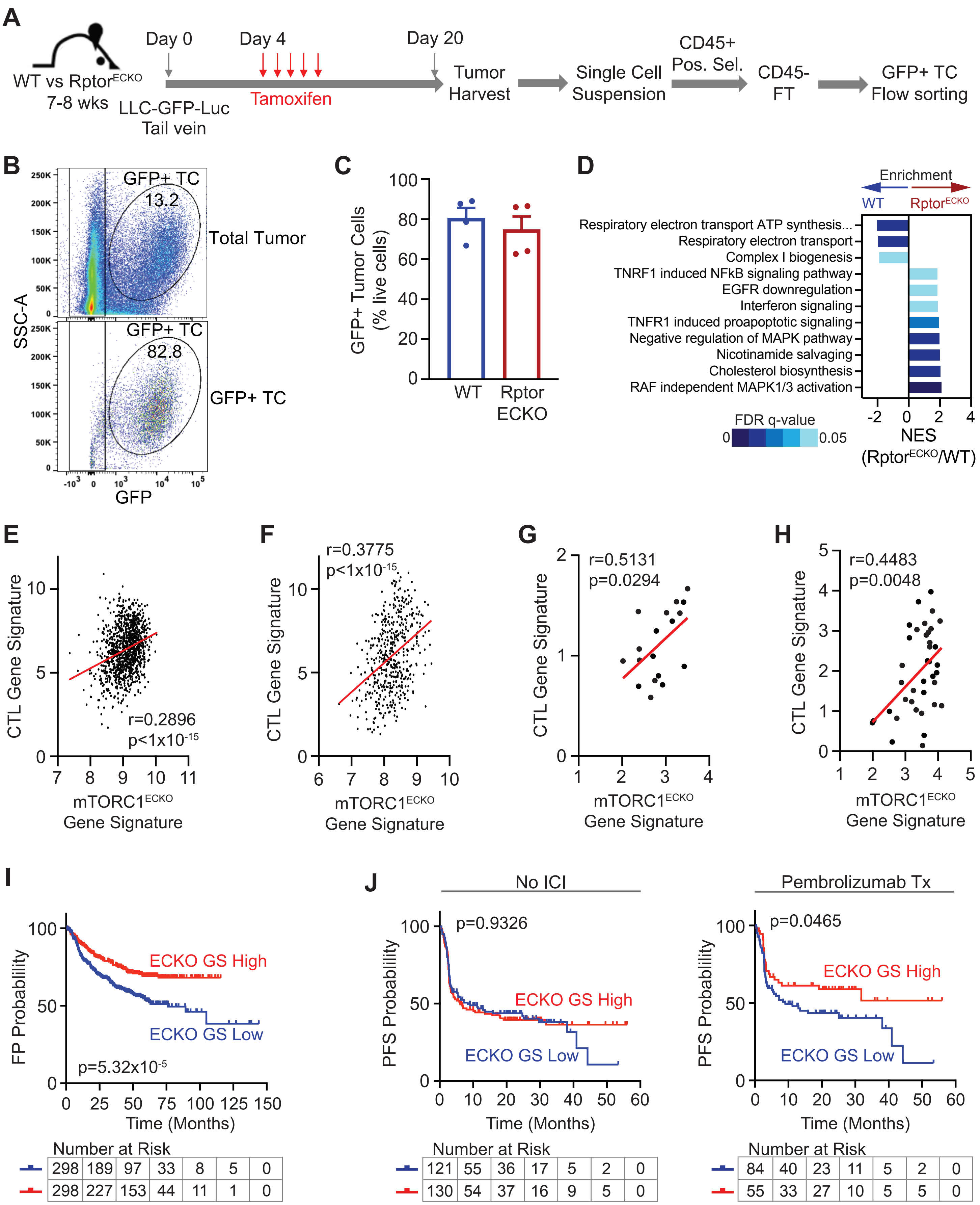
A
WT vs Rptor^ECKO
7-8 wks
Day 0
Day 4
Day 20
LLC-GFP-Luc
Tail vein
Tamoxifen
Tumor Harvest
Single Cell Suspension
CD45+ Pos. Sel.
CD45- FT
GFP+ TC Flow sorting
B
SSC-A
GFP+ TC
13.2
Total Tumor
GFP+ TC
82.8
GFP+ TC
GFP
C
GFP+ Tumor Cells (% live cells)
WT
Rptor ECKO
D
Enrichment
WT
Rptor^ECKO
Respiratory electron transport ATP synthesis...
Respiratory electron transport
Complex I biogenesis
TNRF1 induced NFkB signaling pathway
EGFR downregulation
Interferon signaling
TNFR1 induced proapoptotic signaling
Negative regulation of MAPK pathway
Nicotinamide salvaging
Cholesterol biosynthesis
RAF independent MAPK1/3 activation
FDR q-value
0
0.05
NES (Rptor^ECKO/WT)
E
CTL Gene Signature
r=0.2896
p<1x10^-15
mTORC1^ECKO Gene Signature
F
r=0.3775
p<1x10^-15
CTL Gene Signature
mTORC1^ECKO Gene Signature
G
r=0.5131
p=0.0294
CTL Gene Signature
mTORC1^ECKO Gene Signature
H
r=0.4483
p=0.0048
CTL Gene Signature
mTORC1^ECKO Gene Signature
I
FP Probablility
ECKO GS High
ECKO GS Low
p=5.32x10^-5
Time (Months)
Number at Risk
298 189 97 33 8 5 0
298 227 153 44 11 1 0
J
No ICI
PFS Probablility
p=0.9326
ECKO GS High
ECKO GS Low
Time (Months)
Number at Risk
121 55 36 17 5 2 0
130 54 37 16 9 5 0
Pembrolizumab Tx
PFS Probablility
p=0.0465
ECKO GS High
ECKO GS Low
Time (Months)
Number at Risk
84 40 23 11 5 2 0
55 33 27 10 5 5 0

**Supplemental Figure 8. Human tumors with a high mTORC1$^{ECKO}$ gene signature correlate with improved anti-tumor T cell activity and improved progression-free survival. (A)** Schematic of GFP+ tumor cell sorting of WT and Rptor$^{ECKO}$ lung metastatic tumors. **(B-C)** Flow cytometry confirmed enrichment of GFP+ tumor cells (TC) that is similar between WT and Rptor$^{ECKO}$ tumors. Unpaired t-test, p=0.678. **(D)** Sorted tumor cells from WT or Rptor$^{ECKO}$ lung metastatic tumors were analyzed by RNA-seq and differentially expressed genes were identified. GSEA was performed to identify pathway enrichment. Normalized enrichment score (NES) and false discovery rate (FDR) q-values are indicated. **(E-H)** Differentially expressed genes identified in (D) were used to generate an mTORC1$^{ECKO}$ gene signature. Correlations between the mTORC1$^{ECKO}$ gene signature and a cytotoxic lymphocyte (CTL) gene signature in (E) TCGA LUNG, (F) TCGA SKCM, (G) the Metastatic Breast Cancer Project, and (H) Metastatic Melanoma datasets. **(I)** First progression (FP) survival of lung cancer patients, stratified by low (blue) or high (red) mTORC1$^{ECKO}$ gene signature (ECKO GS). Hazard ratio (HR) is 0.5726. **(J)** Progression-free survival (PFS) of melanoma patients treated with no immune checkpoint inhibitors (ICI, left) or pembrolizumab therapy (right). Patients were stratified by low (blue) or high (red) mTORC1$^{ECKO}$ gene signature (ECKO GS). Hazard ratios (HR) for No ICI and Pembrolizumab therapy is 1.014 and 0.6101, respectively.

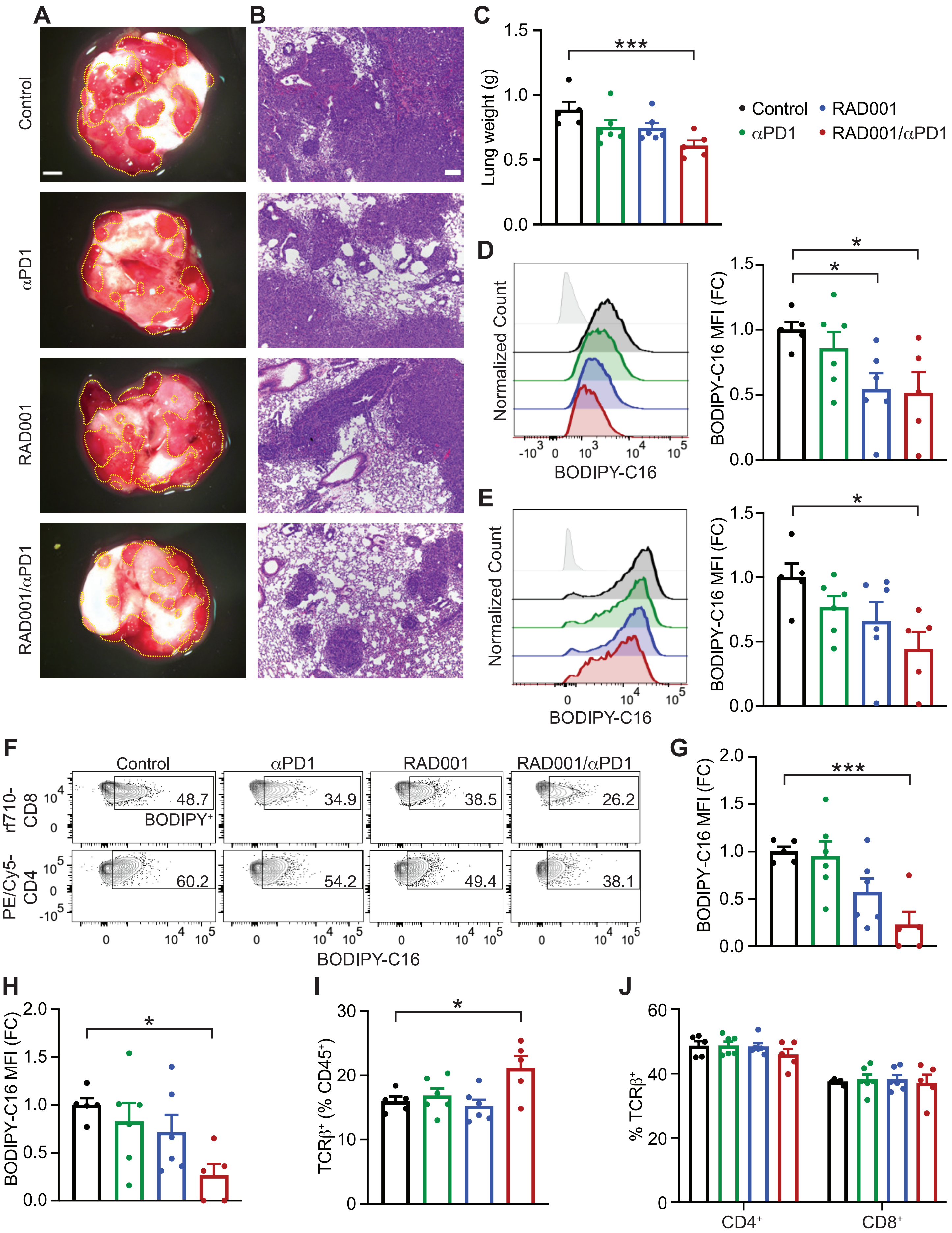

A
B
C
D
E
F
G
H
I
J
Control
αPD1
RAD001
RAD001/αPD1
Lung weight (g)
***
Normalized Count
BODIPY-C16
BODIPY-C16 MFI (FC)
*
rf710-CD8
PE/Cy5-CD4
BODIPY⁺
48.7
34.9
38.5
26.2
60.2
54.2
49.4
38.1
TCRβ⁺ (% CD45⁺)
% TCRβ⁺
CD4⁺
CD8⁺

**Extended Data Figure 9. Low-dose RAD001 combined with anti-PD1 reduces intratumoral BODIPY-C16 transport in lung metastatic tumors.** Female wild type mice were inoculated with E0771-luc cells, as described in Figure 5. **(A)** Representative lung images show reduced surface tumors in RAD001/$\alpha$PD1 animals. Scale bar is 5 mm. Visible tumor area is outlined by the yellow line. **(B)** Representative images H&E staining indicates reduced micro-metastatic tumor burden in RAD001/$\alpha$PD1 lungs. Scale bar is 100 $\mu$m. **(C)** Lung weights were recorded at harvest in grams (g). One-way ANOVA (p=0.0167) with Dunnett's post hoc. **(D)** . BODIPY-C16 median fluorescence intensity (MFI) was determined by flow cytometry in (A) $CD45^{-}CD31^{+}$ endothelial cells and (B) $CD45^{-}CD31^{-}FSC^{hi}$ tumor-cell enriched populations and normalized to Control. Representative histograms are shown. One-way ANOVA ($CD31^{+}$, p=0.0372; tumor-cell enriched, p=0.0443) with Dunnett's post hoc. **(F-H)** BODIPY-C16 MFI was determined in (G) $CD8^{+}$ and (H) $CD4^{+}$ T cells, as described in (D-E). Representative plots are shown in (F). One-way ANOVA ($CD8^{+}$, p=0.00327; $CD4^{+}$, p=0.0367) with Dunnett's post hoc. **(I-J)** T cell populations were determined from treated lung metastatic tumors. (I) T cells ($TCR\beta^{+}$) and (J) $CD4^{+}/CD8^{+}$ T cells are shown as percentage (%) of $CD45^{+}$ or $TCR\beta^{+}$, respectively. Unpaired t-test: $TCR\beta^{+}$, p=0.0182; $CD4^{+}$, p=0.459; $CD8^{+}$, p=0.959. *p<0.05, ***p<0.005.

**Extended Data Table 1. Genotyping primers used in study.**

| Primer Name | | Primer Sequence |
|---|---|---|
| iCdh5-Cre | Forward | 5’-TCCTGATGGTGCCTATCCTC-3’ |
| | Reverse | 5’-CGAACCTGGTCGAAATCAGT-3’ |
| Rptor | Forward | 5’-CTCAGTAGTGGTATGTGCTCAG-3’ |
| | Reverse | 5’-GGGTACAGTATGTCAGCACAG-3’ |
| Rictor | Forward | 5’-GAAGTTATTCAGATGGCCCAGC-3’ |
| | Reverse | 5’-ACTGAATATGTTCATGGTTGTG-3’ |
| Tsc2 | Forward | 5’-TGGCAGGACAGAGGGTCATCATGG-3’ |
| | Reverse | 5’-TTCAGAGTCACCTGGCAGGCTCG-3’ |

**Extended Data Table 2. Mouse flow cytometry antibodies used in study.**

| Antibody | Clone | Dilution | Company | Catalogue # |
|---|---|---|---|---|
| APC/Cy7-CD45 | 30-F11 | 1:500 | BD Biosciences | 557659 |
| PerCP/Cy5.5-TCRβ | H57-597 | 1:500 | Tonbo/Cytek | 65-5961 |
| PE/Cy5-CD4 | GK1.5 | 1:500 | Tonbo/Cytek | 55-0041 |
| PE/Dazzle 594-CD4 | GK1.5 | 1:500 | Biolegend | 100455 |
| Red Fluor 710-CD8 | 53-6.7 | 1:400 | Tonbo/Cytek | 80-0081 |
| PE-CD8 | 53-6.7 | 1:400 | BD Biosciences | 553033 |
| V450-CD107a | 1D4B | 1:200 | BD Biosciences | 560648 |
| PE-Lag3 | C9B7W | 1:400 | eBioscience | 12-2231-81 |
| FITC-PD1 | 29F.1A12 | 1:100 | Biolegend | 135213 |
| PE/Cy7-CD25 | PC61.5 | 1:500 | Tonbo/Cytek | 60-0251 |
| PE-CD127 | A7R34 | 1:500 | Tonbo/Cytek | 50-1271 |
| BV421-CD31 | 390 | 1:200 | Biolegend | 102423 |
| PE/Cy7-EpCAM | G8.8 | 1:500 | Biolegend | 118215 |
| PE-GZMB | NGZB | 1:20 | eBioscience | 12-8898-82 |
| V450-IFNγ | XMG1.2 | 1:100 | Tonbo/Cytek | 75-7311 |
| APC-TNFα | MP6-XT22 | 1:50 | eBioscience | 17-7321-82 |
| eFluor 660-FoxP3 | FJK-16s | 1:250 | eBioscience | 50-5773-82 |
| pS6 (Ser235/236) | | 1:100 | Cell Signaling Technology | 2211 |
| AlexaFluor647 anti-rabbit | | 1:200 | Invitrogen | A21244 |
| PE-IgG2α,κ | eBR2a | 1:20 | eBioscience | 12-4321-81 |
| APC-IgG1,κ | eBRG1 | 1:50 | eBioscience | 17-4301-82 |

**Extended Data Table 3. Quantitative RT-PCR primers used in study.**

| Primer Name | | Primer Sequence |
|---|---|---|
| Mm-*Clstn1* | Forward | 5'-GATGCCGTGGTAGTGGATAAG-3' |
| | Reverse | 5'-CCTGGATGGTGAATGTGTAGTC-3' |
| Mm-*Rptor* | Forward | 5'-CCTCTGTCCATATACGACCT-3' |
| | Reverse | 5'-CTGTGCAGTGCAAACTGT-3' |
| Mm-*Actb* | Forward | 5'-AGAGGGAAATCGTGCGTGAC-3' |
| | Reverse | 5'-CAATAGTGATGACCTGGCCGT-3' |

**Extended Data Table 4. Gene sets used in correlation and survival analysis of bulk RNA datasets.**

| Gene Sets | mTORC1$^{ECKO}$ Gene Signature | | CTL Gene Signature | |
|---|---|---|---|---|
| | DEGs from sorted tumor cells | | Modified from Edwards et al. 2021. | |
| Gene Lists | **Gene Name** | **Gene ID** | **Gene Name** | **Gene ID** |
| | *LCN2* | ENSG00000148346 | *CD8A* | ENSG00000153563 |
| | *PADI1* | ENSG00000142623 | *GZMA* | ENSG00000145649 |
| | *LYPD5* | ENSG00000159871 | *GZMB* | ENSG00000100453 |
| | *ARRB1* | ENSG00000137486 | *GZMM* | ENSG00000197540 |
| | *AMPD3* | ENSG00000133805 | *PRF1* | ENSG00000180644 |
| | *FAT2* | ENSG00000086570 | *IFNG* | ENSG00000111537 |
| | *MXD1* | ENSG00000059728 | | |
| | *VEGFA* | ENSG00000112715 | | |
| | *CRYAB* | ENSG00000109846 | | |
| | *DUSP10* | ENSG00000143507 | | |
| | *C3AR1* | ENSG00000171860 | | |
| | *IER2* | ENSG00000160888 | | |
| | *PTGS2* | ENSG00000073756 | | |
| | *SCN8A* | ENSG00000196876 | | |
| | *SGK1* | ENSG00000118515 | | |
| | *ERRFI1* | ENSG00000116285 | | |
| | *GPR35* | ENSG00000178623 | | |
| | *BDNF* | ENSG00000176697 | | |
| | *RCAN1* | ENSG00000159200 | | |
| | *GPR39* | ENSG00000183840 | | |
| | *DUSP1* | ENSG00000120129 | | |
| | *SOCS3* | ENSG00000184557 | | |
| | *TSC22D1* | ENSG00000102804 | | |
| | *ST3GAL1* | ENSG00000008513 | | |
| | *FILIP1L* | ENSG00000168386 | | |
| | *BACH1* | ENSG00000156273 | | |
| | *DNAJB9* | ENSG00000128590 | | |
| | *ARRDC3* | ENSG00000113369 | | |
| | *RBPJ* | ENSG00000168214 | | |
| | *HSPA5* | ENSG00000044574 | | |
| | *BIN1* | ENSG00000136717 | | |
| | *ITGB2* | ENSG00000160255 | | |
| | *TEAD2* | ENSG00000074219 | | |
| | *CGREF1* | ENSG00000138028 | | |
| | *FGFR1* | ENSG00000077782 | | |
| | *RAP1GAP* | ENSG00000076864 | | |
| | *CD109* | ENSG00000156535 | | |
| | *MICAL1* | ENSG00000135596 | | |
| | *PAPLN* | ENSG00000100767 | | |
| | *ACSL6* | ENSG00000164398 | | |
| | *CD22* | ENSG00000012124 | | |

**Extended Table 5. Gene sets used in single-cell RNA-seq analysis of endothelial cells.**

| MsigDB Gene Sets | REACTOME_mTORC1_ mediated_pathway | | REACTOME_RAB_regulation_ of_trafficking | |
|---|---|---|---|---|
| Collection | C2, CP | | C2, CP | |
| Gene Lists | Gene Name | Gene ID | Gene Name | Gene ID |
| | *AKT1S1* | ENSG00000204673 | *AKT1* | ENSG00000142208 |
| | *EEF2K* | ENSG00000103319 | *AKT2* | ENSG00000105221 |
| | *EIF4B* | ENSG00000063046 | *AKT3* | ENSG00000117020 |
| | *EIF4E* | ENSG00000151247 | *ALS2* | ENSG00000003393 |
| | *EIF4EBP1* | ENSG00000187840 | *ALS2CL* | ENSG00000178038 |
| | *EIF4G1* | ENSG00000114867 | *ANKRD27* | ENSG00000105186 |
| | *LAMTOR1* | ENSG00000149357 | *ARF6* | ENSG00000165527 |
| | *LAMTOR2* | ENSG00000116586 | *CCZ1* | ENSG00000122674 |
| | *LAMTOR3* | ENSG00000109270 | *CCZ1B* | ENSG00000146574 |
| | *LAMTOR4* | ENSG00000188186 | *CHM* | ENSG00000188419 |
| | *LAMTOR5* | ENSG00000134248 | *CHML* | ENSG00000203668 |
| | *MLST8* | ENSG00000167965 | *DENND1A* | ENSG00000119522 |
| | *MTOR* | ENSG00000198793 | *DENND1B* | ENSG00000213047 |
| | *RHEB* | ENSG00000106615 | *DENND1C* | ENSG00000205744 |
| | *RPS6* | ENSG00000137154 | *DENND2A* | ENSG00000146966 |
| | *RPS6KB1* | ENSG00000108443 | *DENND2B* | ENSG00000166444 |
| | *RPTOR* | ENSG00000141564 | *DENND2C* | ENSG00000175984 |
| | *RRAGA* | ENSG00000155876 | *DENND2D* | ENSG00000162777 |
| | *RRAGB* | ENSG00000083750 | *DENND3* | ENSG00000105339 |
| | *RRAGD* | ENSG00000025039 | *DENND4A* | ENSG00000174485 |
| | *SLC38A9* | ENSG00000177058 | *DENND4B* | ENSG00000198837 |
| | *YWHAB* | ENSG00000166913 | *DENND4C* | ENSG00000137145 |
| | | | *DENND5A* | ENSG00000184014 |
| | | | *DENND5B* | ENSG00000170456 |
| | | | *DENND6A* | ENSG00000174839 |
| | | | *DENND6B* | ENSG00000205593 |
| | | | *GABARAP* | ENSG00000170296 |
| | | | *GABARAPL2* | ENSG00000034713 |
| | | | *GAPVD1* | ENSG00000165219 |
| | | | *GDI1* | ENSG00000203879 |
| | | | *GDI2* | ENSG00000057608 |
| | | | *GGA1* | ENSG00000100083 |
| | | | *GGA2* | ENSG00000103365 |
| | | | *GGA3* | ENSG00000125447 |
| | | | *HPS1* | ENSG00000107521 |
| | | | *HPS4* | ENSG00000100099 |
| | | | *MADD* | ENSG00000110514 |
| | | | *MAP1LC3B* | ENSG00000140941 |
| | | | *MON1A* | ENSG00000164077 |
| | | | *MON1B* | ENSG00000103111 |
| | | | *OPTN* | ENSG00000123240 |
| | | | *RAB10* | ENSG00000084733 |
| | | | *RAB11A* | ENSG00000103769 |
| | | | *RAB11B* | ENSG00000185236 |
| | | | *RAB12* | ENSG00000206418 |
| | | | *RAB13* | ENSG00000143545 |
| | | | *RAB14* | ENSG00000119396 |
| | | | *RAB18* | ENSG00000099246 |

| | | | | |
|---|---|---|---|---|
| | | | *RAB1A* | ENSG00000138069 |
| | | | *RAB1B* | ENSG00000174903 |
| | | | *RAB21* | ENSG00000080371 |
| | | | *RAB27A* | ENSG00000069974 |
| | | | *RAB27B* | ENSG00000041353 |
| | | | *RAB31* | ENSG00000168461 |
| | | | *RAB32* | ENSG00000118508 |
| | | | *RAB33A* | ENSG00000134594 |
| | | | *RAB33B* | ENSG00000172007 |
| | | | *RAB35* | ENSG00000111737 |
| | | | *RAB38* | ENSG00000123892 |
| | | | *RAB39A* | ENSG00000179331 |
| | | | *RAB39B* | ENSG00000155961 |
| | | | *RAB3A* | ENSG00000105649 |
| | | | *RAB3GAP1* | ENSG00000115839 |
| | | | *RAB3GAP2* | ENSG00000118873 |
| | | | *RAB3IL1* | ENSG00000167994 |
| | | | *RAB3IP* | ENSG00000127328 |
| | | | *RAB4A* | ENSG00000168118 |
| | | | *RAB5A* | ENSG00000144566 |
| | | | *RAB5B* | ENSG00000111540 |
| | | | *RAB5C* | ENSG00000108774 |
| | | | *RAB6A* | ENSG00000175582 |
| | | | *RAB6B* | ENSG00000154917 |
| | | | *RAB7A* | ENSG00000075785 |
| | | | *RAB7B* | ENSG00000276600 |
| **Gene Lists (cont.)** | | | *RAB8A* | ENSG00000167461 |
| | | | *RAB8B* | ENSG00000166128 |
| | | | *RAB9A* | ENSG00000123595 |
| | | | *RABEP1* | ENSG00000029725 |
| | | | *RABGAP1* | ENSG00000011454 |
| | | | *RABGEF1* | ENSG00000154710 |
| | | | *RGP1* | ENSG00000107185 |
| | | | *RIC1* | ENSG00000107036 |
| | | | *RIN1* | ENSG00000174791 |
| | | | *RIN2* | ENSG00000132669 |
| | | | *RIN3* | ENSG00000100599 |
| | | | *RINL* | ENSG00000187994 |
| | | | *SBF1* | ENSG00000100241 |
| | | | *SBF2* | ENSG00000133812 |
| | | | *SYTL1* | ENSG00000142765 |
| | | | *TBC1D10A* | ENSG00000099992 |
| | | | *TBC1D10B* | ENSG00000169221 |
| | | | *TBC1D10C* | ENSG00000175463 |
| | | | *TBC1D13* | ENSG00000107021 |
| | | | *TBC1D14* | ENSG00000132405 |
| | | | *TBC1D15* | ENSG00000121749 |
| | | | *TBC1D16* | ENSG00000167291 |
| | | | *TBC1D17* | ENSG00000104946 |
| | | | *TBC1D2* | ENSG00000095383 |
| | | | *TBC1D20* | ENSG00000125875 |
| | | | *TBC1D24* | ENSG00000162065 |
| | | | *TBC1D25* | ENSG00000068354 |

| | | | | |
|---|---|---|---|---|
| | | | *TBC1D3* | ENSG00000274611 |
| **Gene Lists (cont.)** | | | *TBC1D7* | ENSG00000145979 |
| | | | *TRAPPC1* | ENSG00000170043 |
| | | | *TRAPPC10* | ENSG00000160218 |
| | | | *TRAPPC11* | ENSG00000168538 |
| | | | *TRAPPC12* | ENSG00000171853 |
| | | | *TRAPPC13* | ENSG00000113597 |
| | | | *TRAPPC2* | ENSG00000196459 |
| | | | *TRAPPC2L* | ENSG00000167515 |
| | | | *TRAPPC3* | ENSG00000054116 |
| | | | *TRAPPC4* | ENSG00000196655 |
| | | | *TRAPPC5* | ENSG00000181029 |
| | | | *TRAPPC6A* | ENSG00000007255 |
| | | | *TRAPPC6B* | ENSG00000182400 |
| | | | *TRAPPC8* | ENSG00000153339 |
| | | | *TRAPPC9* | ENSG00000167632 |
| | | | *TSC1* | ENSG00000165699 |
| | | | *TSC2* | ENSG00000103197 |
| | | | *ULK1* | ENSG00000177169 |
| | | | *YWHAE* | ENSG00000108953 |
